# CLONALITY AND POLYPLOIDY CONTRIBUTE TO THE SPREAD OF *AVRAINVILLEA LACERATA* (BRYOPSIDALES, CHLOROPHYTA) IN HAWAIʻI

**DOI:** 10.1101/2024.05.04.592553

**Authors:** Brinkley M. Thornton, Heather L. Spalding, Solenn Stoeckel, Melissa L. Harris, Rachael M. Wade, Stacy A. Krueger-Hadfield

**Author notes:** California State Polytechnic University Humboldt, Arcata, CA, 95521, USA Running head: Clonality and polyploidy in *Avrainvillea. Corresponding author: Stacy A. Krueger-Hadfield, Virginia Institute of Marine Science Eastern Shore Laboratory.

## Abstract

The relative rates of sexual versus asexual reproduction influence the partitioning of genetic diversity within and among populations. During range expansions, uniparental reproduction often facilitates colonization and establishment. The arrival of the green alga *Avrainvillea lacerata* has caused shifts in habitat structure and community assemblages since its discovery in 1981 offshore of west Oʻahu, Hawai‘i. Field observations suggest this species is spreading via vegetative reproduction. To characterize the reproductive system of *A. lacerata* in Hawai‘i, we developed seven microsatellite loci and genotyped 321 blades collected between 2018 and 2023 from two intertidal sites at Maunalua Bay and ʻEwa Beach. We found one to four alleles at multiple loci, suggesting *A. lacerata* is tetraploid. Each site was characterized by high genotypic richness (R > 0.8). However, clonal rates were also high at both sites, suggesting vegetative spread of *A. lacerata* plays a significant role. The importance of clonal reproduction for the persistence of *A. lacerata* in Hawai‘i is consistent with the ecological data collected for this species, and observations of other abundant macroalgal invaders in Hawaiʻi and other regions of the world. These data demonstrate the necessity for implementing appropriate population genetic methods and provide insights into the biology of this alga that will contribute to future studies on effective management strategies incorporating its reproductive system. This study represents one of the few investigating green algal population genetic patterns and contributes to our understanding of algal reproductive system evolution.

## INTRODUCTION

The reproductive system describes the relative rates of sexual and asexual reproduction, governing genetic diversity through the transmission of genes from one generation to the next (Barrett, 2011). Sexual reproduction ranges from self-fertilization (also called selfing) to outcrossing, whereby outcrossing confers greater standing genetic diversity due to cross-fertilization between individuals (Hamrick & Godt, 1996). Selfing, the union of gametes produced by the same individual, lowers genetic diversity through reduced effective recombination and increased levels of homozygosity (Charlesworth & Charlesworth, 1987; Charlesworth & Wright, 2001; Ellstrand & Elam, 1993). Offspring produced by most asexual processes (or clonality) are genetic copies of their parent, barring mutation (de Meeûs et al., 2007; Orive & Krueger-Hadfield, 2021). Contrary to selfing, asexual reproduction tends to favor the maintenance of parental heterozygosity, and may even increase heterozygosity via the accumulation of mutations over time (Marshall & Weir, 1979; Reichel et al., 2016; Stoeckel & Masson, 2014). The relative rates of sexual versus asexual reproduction are therefore integral to our understanding of population structure. Yet, most of our understanding of the variation in reproductive modes among populations is based on angiosperms (e.g., Whitehead et al., 2018) or animals (e.g., Olsen et al., 2020), and primarily focuses on sexual reproduction (Lane et al., 2011; Rushworth et al., 2022).

Traits that influence the reproductive system can rapidly shift in response to changes in environment or demography (Barrett, 2008; Eckert et al., 2010). Biological invasions are excellent examples of anthropogenically mediated range expansions (Pannell et al., 2015) and are among the main threats to biodiversity (Kolar & Lodge, 2001; Bellard et al., 2016; Dueñas et al., 2021). Following an invasion, individuals in small, recently established populations often exhibit shifts to uniparental reproduction (i.e., selfing, asexual reproduction, or some combination of both; Pannell et al., 2015). There are often only a few founding individuals, limiting mating opportunities. Therefore, uniparental reproduction provides reproductive assurance (i.e., Baker’s Law; Baker, 1955; Pannell, 2015). Patterns of geographic parthenogenesis (originally described by Vandel, 1928; see also Bierzychudek, 1985; Lynch, 1984) have been extended to invasions (Pannell et al., 2015) whereby asexual reproduction can maintain high fitness genotypes that otherwise could be lost in small populations through sexual reproduction and genetic drift (Haag & Ebert, 2004). Moreover, asexual reproduction favors population growth as mating is prevented between poorly adapted individuals (Antonovics, 1968; Peck et al., 1998). Yet, most population genetic theory has been developed for obligate sexual reproduction and we lack the appropriate tools to accurately assess and interpret rates of asexual reproduction in natural populations (see Arnaud-Haond et al., 2007; Stoeckel et al., 2021).

The lack of population genetic data is particularly acute for macroalgae despite these taxa constituting a large portion of non-native species in marine environments (Schaffelke et al., 2006; Williams & Smith, 2007). Macroalgal invasions have concomitant negative consequences for local communities via the overgrowth and smothering of native species, alteration of surrounding habitats, and a decrease in available food sources for herbivores (Schaffelke et al., 2006). Despite the continuous increase in the number of macroalgal invasions, most of the knowledge on invasive macroalgae is limited to what we have learned from high-profile invaders, such as *Caulerpa taxifolia* (Deveney et al., 2008) and *Sargassum muticum* (Engelen et al., 2015), and mainly focused on ecological rather than evolutionary processes (see discussion in Sotka et al., 2018), especially with regard to reproductive system variation (see Krueger-Hadfield et al., 2021; Krueger-Hadfield, 2020). Algae display tremendous variation in life cycles (Bell, 1994) and are predicted to be partially clonal (i.e., undergo sexual and asexual reproduction simultaneously; Otto & Marks, 1996). Outcrossing should favor the diploid life cycle while inbreeding and asexual reproduction should favor prolonged haploid phases in the life cycle (Otto & Marks, 1996). Yet, there were too few data to test this hypothesis and this has not improved substantially in the past ∼30 years (see as an example, Krueger-Hadfield et al., 2021; Krueger-Hadfield, 2024).

Here, we investigate the reproductive mode of the green alga *Avrainvillea lacerata* J.Agardh. In 1981, *Avrainvillea*, morphologically identified as *A. amadelpha,* was documented in Hawaiʻi off the western shores of Oʻahu (Brostoff, 1989). Nearly 40 years later, molecular assessments with the inclusion of type material revealed this alga to be *A. lacerata* (Lagourgue et al., 2023; Wade, 2019). Since the initial discovery, new observations of *A. lacerata* have been documented around Oʻahu, both in the intertidal as well as the mesophotic (Cox et al., 2017; Spalding et al., 2019). This alga is now found on the neighboring islands of Kaua‘i and Maui (Smith et al., 2002; Wade, 2019). The invasion of *A. lacerata* has led to habitat and community change because it forms extensive mounds, some up to 30 meters wide (Littler et al., 2004, 2005; Peyton, 2009). The mounds alter the benthos through increased sedimentation, which modifies hard substrate to resemble soft sediment habitats (Foster et al., 2019; Littler et al., 2004). The alga’s ability to engineer habitat structure, complex branching and holdfast morphology (Littler & Littler, 1992; Olsen-Stojkovich, 1985), and possible herbivore-deterring secondary metabolites (see Hay et al., 1990) have all influenced the surrounding ecosystem by contributing towards significant shifts in surrounding invertebrate (Foster et al., 2019; Longenecker et al., 2011), algal (Peyton, 2009; Smith et al., 2002), and fish communities (Langston & Spalding, 2017).

While the ecological consequences of the *Avrainvillea lacerata* invasion on Hawaiian ecosystems have been documented (e.g., Foster et al., 2019; Peyton, 2009; Smith et al., 2002; Veazey et al., 2019), there is a lack of basic knowledge regarding the reproductive system of both this species and the genus more broadly. For example, the life cycle is assumed to be diploid, in which somatic development only occurs in the diploid phase. However, this assumption is based on *A. lacerata*’s phylogenetic position relative to other green algae in the Bryopsidales and in the genus *Avrainvillea* (Littler & Littler, 1992; Olsen-Stojkovich, 1985). Polyploidy has been suggested for some bryopsidalean macroalgae (Kapraun, 1994; Kapraun & Nguyen, 1994; Kapraun & Shipley, 1990; Varela-Álvarez et al., 2012), but no studies have investigated the extent of polyploidy throughout this order using genetic tools. Moreover, the life cycle influences the reproductive system (see Otto and Marks 1996, Krueger-Hadfield 2020), but there are no population genetic data describing the prevailing reproductive mode or overall reproduction in the genus *Avrainvillea* (Littler & Littler, 1992; Olsen-Stojkovich, 1985). Field observations suggest the alga is predominately spreading through the vegetative growth of mounds, and reproductive structures have never been observed (Smith et al., 2002). To our knowledge, no detailed phenology study exists to complement studies of reproduction in *A. lacerata*.

In this study, we developed seven microsatellite loci to genotype *Avrainvillea lacerata* blades and describe its reproductive system in Hawaiʻi. Based on field observations of massive mounds suggesting vegetative spread and the lack of reproductive thalli, we hypothesized that *A. lacerata* is primarily spreading through asexual reproduction. We considered asexual reproduction as the only uniparental reproductive mode because *A. lacerata* is hypothesized to be dioecious based on its phylogenetic position (Olsen-Stojkovich, 1985; Verbruggen et al., 2009). As such, selfing would not be possible, though inbreeding can still occur between related individuals. Further, we sampled over multiple years to quantify the clonal rate at the sites sampled (see discussion in Becheler et al., 2017). These microsatellites will facilitate future studies investigating the invasion history of this alga and could eventually discern the source of the invasion from its native range (see as example Krueger-Hadfield et al., 2017). Not only will quantitative data on the reproductive system aid in the management of this invader, it will test the Otto and Marks (1996) hypothesis of a correlation between the reproductive mode and life cycle, expanding the available population genetic data for macroalgae across marine biomes.

## METHODS

### Field sampling

We collected *Avrainvillea lacerata* blades from two sites along the southern coastline of Oʻahu. Single blades were considered a putative genet (i.e., one genotype, Harper, 1980), with a blade defined as a single upright fan-shaped apex connected to a single stipe from a clump with a discrete holdfast mound (i.e., a ramet). At each site, we haphazardly placed a 50 meter transect parallel to the shore during a low tide in a representative area of the intertidal zone occupied by *A. lacerata*. At each sampling point, we collected a single upright blade from distinct mounds separated by at least one meter. In May 2018 and June 2019, we collected 50 blades from Maunalua Bay (21.279880, -157.728160) and 50 blades from the ʻEwa Beach site, Lagoon East (21.304310, -158.036240; site 13, Cox et al., 2017) (Figure 1).

**Figure 1.**
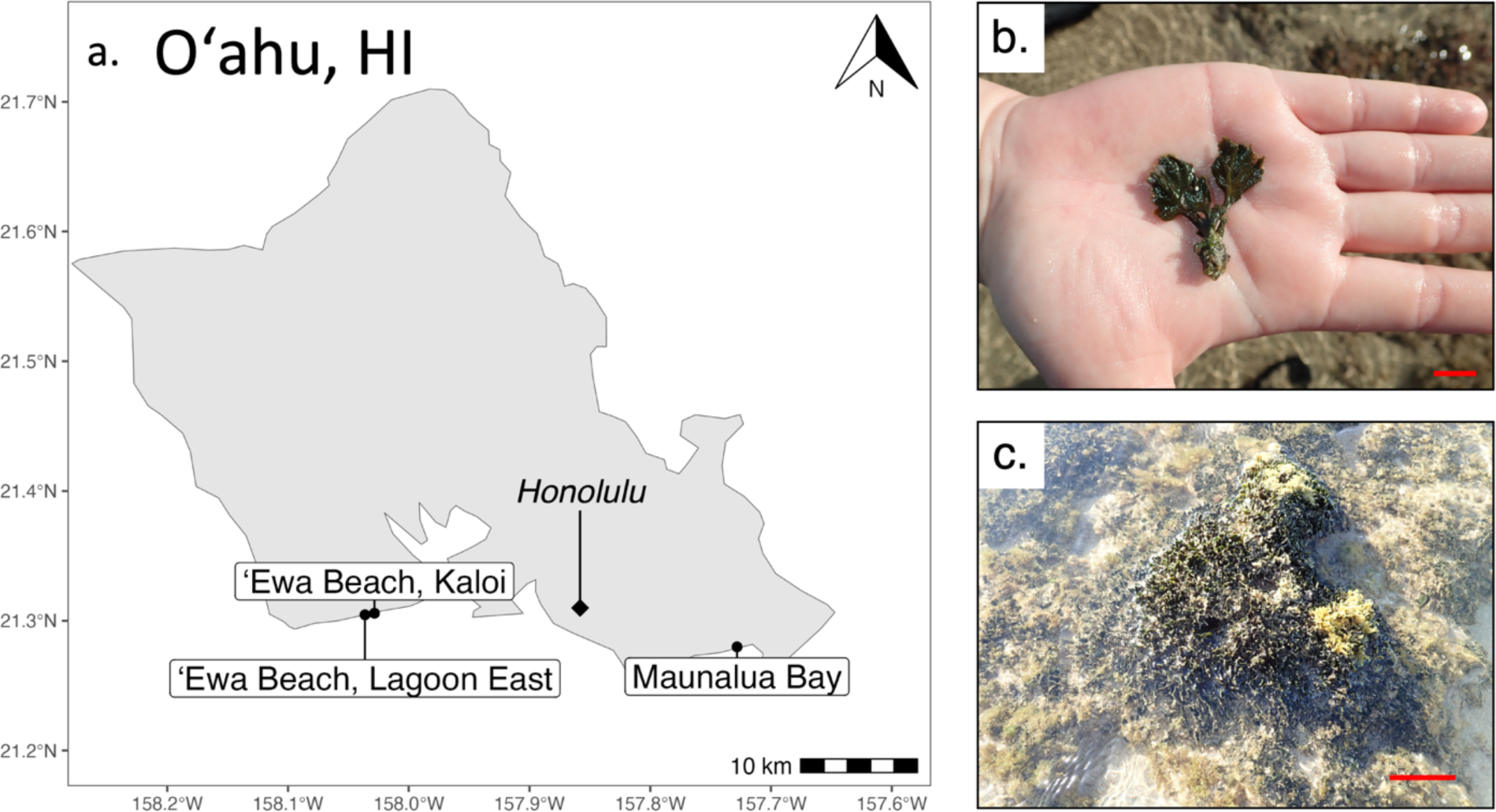
Sampling sites of *Avrainvillea lacerata* on Oʻahu, HI. (a) Map of *A. lacerata* sampling sites on Oʻahu, HI. Honolulu is included as a locational reference. (b) One thallus of *A. lacerata* with multiple blades connected to a single holdfast. Scalebar: 1 cm. Photo credit: H. Spalding. (c) Example of an *A*. *lacerata* mound from which blades were sampled. Scalebar: 10 cm. Photo credit: H. Spalding.

Our site in Maunalua Bay is a protected and the substrate is comoposed of soft sediment. Increased coastal development has led to increased sedimentation (Peyton, 2009). There has also been an increase in invasive species, such as *Avrainvillea lacerata* in this area (Peyton, 2009). Concomitant with all the changes, there has been a decrease in the native seagrass *Halophila hawaiiana* (Peyton, 2009). In 1985, *A. lacerata* was collected in Maunalua Bay (Brostoff, 1989). By 2003, there were extensive mounds that covered up to 100% of the soft sediment (Peyton, 2009). Despite a large scale removal effort, mounds of *A. lacerata* have grown back, though in a much patchier distribution than before the removal. We predict the occurrence of more unique genotypes at our site in Maunalua Bay as compared to the site ‘Ewa (see below) from (i) regrowth of mounds that were left behind following the removal, (ii) the arrival of new genotypes from sexual reproduction, or (iii) the arrival of new genotypes from nearby sites, including the subtidal or mesophotic.

‘Ewa Beach Lagoon East by comparison is a more exposed site with greater wave exposure and hard substratum. This site also has greater native algal diversity (see Cox et al., 2017). In 2008, *Avrainvillea lacerata* mounds were found in low abundance (Cox et al., 2017), but now it is one of the most abundant algae, forming extensive mounds. Due to greater wave action and the competition for space, we predict recruitment of *A. lacerata* will be more difficult as compared to the Maunalua Bay site. Thus, we expected to find a higher occurrence of vegetative growth of mounds leading to the dominance of fewer genotypes.

In addition to comparing population genetic structure between the two sites at Maunalua Bay and ʻEwa Beach, we wanted to make comparisons between two sites that were ∼870 meters apart but differed in their environmental characteristics. In May 2021, at ʻEwa Beach, we collected 30 blades each from Lagoon East and a second site called Kaloi (21.305800, - 158.028400; site 2, Cox et al., 2017) (Figure 1). Kaloi differs from Lagoon East in substrate angularity and topographic relief and is a flat bench with higher wave exposure. Lagoon East, by contrast, is more wave protected in comparison to Kaloi and has an angular carbonate bench with greater topographic relief.

In addition to the population-level sampling, we wanted to assess diversity within single *Avrainvillea lacerata* mounds, which result from lateral vegetative growth (Littler et al., 2005), presumably of a single genet. In May 2022, we collected blades from eight mounds at Lagoon East. From each mound, we collected five blades for a total of 40 blades. We repeated this sampling in June 2023, in which we sampled blades from eight separate mounds in ʻEwa, Lagoon East. Each clump had between 1-3 holdfasts fused together where each holdfast seemed to be associated with a group of blades. From each holdfast within a clump, 2-3 blades were sampled.

All blades were rinsed in seawater and preserved in silica gel (ACTÍVA Flower Drying Art Silica Gel, Marshall, TX, U.S.A).

### DNA extraction

We extracted DNA from approximately 5-6 mg from each dried *Avrainvillea lacerata* blade. We homogenized blades with one 2.8 mm ceramic bead using a bead mill (BeadMill24, Fisher Scientific, Waltham, MA, USA) at a speed of 4 m/sec for 45 seconds until each blade was homogenized into a fine powder. Some blades required up to four bursts with the bead mill. We used the OMEGA biotek E.Z.N.A.^®^ Plant DNA DS Kit (Norcross, GA, U.S.A.) according to the manufacturer’s instructions. We eluted DNA in a single volume of 150 μL using the provided elution buffer.

### Microsatellite locus identification

Single sequence repeat (SSR)-enriched genomic sequence data was generated by Microsynth ecogenics GmbH (Balgach, Switzerland). We identified putative loci from the SSR-enriched library and followed Schoebel et al. (2013), with modifications implemented in Ryan et al. (2021) and Heiser et al. (2023) which we summarize here. We used MSATCOMMANDER 1.0.8-beta (Faircloth, 2008) to design primers for di-, tri- and tetranucleotide repeat motifs separately. A minimum of eight repeats was selected and the following primer melting temperatures (T_m_): minimum of 58°C, optimum of 60°C, and maximum of 62°C. For dinucleotides, we identified 408 sequences with eight or more repeats, 95 of those had primers assigned, and 16 were potentially duplicated in the library. For trinucleotides, we identified 1712 sequences with eight or more repeats, 376 of those had primers assigned, and 102 were potentially duplicated in the library. For tetranucleotides, we identified 153 sequences with six or more repeats, 22 of those had primers assigned, and two were potentially duplicated in the library. We had 79, 274, and 20 potential loci with di-, tri-, and tetranucleotide repeat motifs, respectively.

We followed Schoebel et al. (2013) using R version 3.5.2 (R Core Team, 2018) to combine the primer and microsatellite sequences into one file. For the dinucleotides, we had 76 unique reads remaining after merging the files. For trinucleotides, we had 267 unique reads remaining after merging the files. For tetranucleotides, we had 20 unique reads remaining after merging the files. After removing duplicated forward and reverse primer sequences, we had 239 unique reads remaining. We then combined the files with unique reads.

We calculated the absolute difference between the forward and reverse T_m_ for each primer pair and sorted from smallest (0°C) to largest (2.51°C). We, then, sorted the putative loci by the forward penalty score, reverse penalty score, and the pair penalty score. Lastly, we calculated and sorted by the absolute difference between penalty scores divided by the pair penalty from smallest to largest to ensure that the difference between the forward and reverse penalties was as small as possible. We selected the top 162 loci from each of those five categories and combined them in one file. We ranked the resulting 94 loci through the combined score from all five categories.

Finally, we conducted a BLAST search in Geneious Prime v.2022.2.2 (Biomatters, Ltd., Auckland, New Zealand) using the SSR-enriched library to ensure that only one primer pair was binding to the same locus, no primer pair was binding to more than one locus, and repeat regions were not within the primers. A total of 49 candidate loci were initially screened using seven blades (see Table S1).

### Microsatellite locus screening, PCR conditions, and fragment analysis

Candidate loci were amplified using simplex PCRs with a final volume 15 µL: 2 μL of diluted DNA (1:100), 250 nM of each forward and reverse oligo, 1X of GoTaq® Flexi DNA Green Buffer (Promega, Cat # M891A), 2 mM of MgCl_2_, 250 μM of each dNTP (Promega, Cat # R0192), 1mg/mL of bovine serum albumin (BSA, Fisher Bioreagents, Cat # BP9706-100i), and 1U of Promega GoTaq® Flexi DNA Polymerase. We used the following PCR program: 95°C for two minutes, followed by 30 cycles of 95°C for 30 seconds, 56°C for 30 seconds, and 72°C for 30 seconds, with a final elongation at 72°C for five minutes. Approximately 5 µL of each PCR product was screened on 1.5% agarose gels stained with GelRed (Biotium, Fremont, CA, USA, Cat #41002-1). Each locus was then categorized based on the amplification profile: one band, multiple bands, or no amplification (Table S1). We considered loci to be good candidates if they amplified well across the seven blades and only had one band per thallus. Based on these criteria, 12 candidate loci were selected for screening using the capillary sequencer.

We assigned dyes – 6FAM, NED, VIC, PET – to each forward oligo for the 12 loci (Table S1). We performed fragment analysis of all samples at the Heflin Center for Genomic Sciences at the University of Alabama at Birmingham. We diluted 1 μL of PCR product in 10 µL of molecular grade H_2_O and then added 1.5 μL diluted PCR product to 9.7 μL HiDi formamide (Applied Biosystems) and 0.35 μL GS500 LIZ (Applied Biosystems, Cat # 4322682).

We scored alleles using GENEIOUS PRIME v.2022.2.2 (https://www.geneious.com). We implemented strict criteria in scoring alleles in which we scored alleles that (a) exhibited 500 or more relative fluorescence units (RFU) and (b) had a peak architecture common to all other alleles for a particular locus. For each allele we scored, we recorded the RFU in addition to the base pair size. Following this, we binned raw allele sizes manually into integers, ensuring rounding did not artificially increase allelic diversity. A considerable proportion of blades displayed more than two alleles in at least one of the loci studied. To confirm this, we ran re-extracted DNA from a subset of blades and repeated PCRs.

### Flow Cytometry

We used lab-reared *Arabidopsis thaliana* to establish a working protocol. We released nuclei from *A. thaliana* by finely chopping freshly grown roots with a sterile razor blade in 1 mL of LB01 buffer (Doležel et al., 1989) until homogenized (∼two minutes). We incubated the nuclear suspension at 4°C on a 3D Platform Rotator (Fisher Scientific, Hampton, NH) for 15 minutes. The nuclear suspension was filtered through a pre-wet (with 500 μL of buffer) 40 μm nylon mesh cell strainer to remove large fragments and subsequently added 50 μg/mL of propidium iodide (PI) to stain the nuclei. Following a five-minute incubation period in the dark, we analyzed the fluorescence intensity of the nuclei using an S3e Cell Sorter (BIO-RAD, Hercules, CA). The cell sorter excited the PI with a 488 nm laser and detected its emission with a 615/25 nm band pass filter. We recorded and analyzed the intensities of the fluorescence emitted by the isolated nuclei using ProSort (BIO-RAD, Hercules, CA). Once we received live *Avrainvillea lacerata* blades, we repeated the same protocol. We removed residual epiphytes or sediment by rinsing *A. lacerata* blades with sterile artificial seawater (Harrison et al., 1980). Next, we released nuclei from *A. lacerata* by cutting approximately 1-2 cm^2^ of a blade with dissecting scissors in 1 mL of the LB01 buffer and following the same methods described above for *Arabidopsis*. Flow cytometry data was analyzed using FlowJo^TM^ ver. 10.8.1 (Becton, Dickinson and Company, Ashland, OR).

### Fluorescence Microscopy

We rehydrated silica-dried *Avrainvillea lacerata* blades in sterile artificial seawater (Harrison et al., 1980) overnight at room temperature (∼20°C). For each blade, we placed a small piece (∼2-3 mm^2^) of the rehydrated blade into a glass petri dish. Using a 5% Dawn dish soap solution (Procter & Gamble, Cincinnati, OH) and dissection needles, we teased apart individual siphons under a dissection scope. Following this, we transferred the siphons to slides pretreated with subbing solution. Finally, we simultaneously mounted and stained the siphons with Fluoromount-G mounting medium with DAPI (Invitrogen, Waltham, MA). We captured DAPI images of nuclei with a CoolSNAP MYO CCD camera (Photometrics, Tuscan, AZ) on a Nikon Eclipse 80i fluorescence microscope.

### Assigning polyploid genotypes

We observed between one and four alleles at each locus per blade (Table S1). We could easily assign a tetraploid genotype for blades with a single allele (e.g., AAAA) or four alleles observed at a locus (e.g., ABCD). However, for blades with two or three alleles at a single locus, we first determined allele dosage. GenoDive (Meirmans, 2020) was used to correct for unknown allele dosage through imputation of missing alleles using maximum likelihood based on estimated allele frequencies observed in the genotyped specimens prior to allele dosage estimation (see Preston et al., 2022). As we obtained greater genotypic diversity than expected based on raw alleles observed from blades collected from the same mound (see Results), we developed a method whereby we used a multinomial kernel to compute the likelihood of each possible genotype and used the RFU for each allele as a proxy for the “dosage”. For each blade, we assigned the genotype that corresponded with the maximum posterior probability.

### Population genetic analyses

We calculated the following summary statistics at each site and time point using GenAPoPop, a software to analyze genetic diversity and reproductive mode in diploid and autopolyploid populations (Stoeckel et al., 2024). First, we calculated genotypic richness (*R*) using the formula 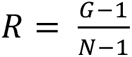, where *G* is the number of unique multilocus genotypes (MLGs) and *N* is the number of blades genotyped (Dorken & Eckert, 2001). Next, we assessed genotypic evenness using *pareto β* (see Box 4, equation 17, Arnaud-Haond et al., 2007) and *D*\* (see Box 3, equation 11, Arnaud-Haond et al., 2007). We, then, estimated linkage disequilibrium (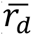, Agapow & Burt, 2001). We calculated expected heterozygosity (*H_E_,* Nei, 1978), observed heterozygosity (*H_O_*, Hardy, 2016), and the inbreeding coefficient (*F_IS_*, Weir & Cockerham, 1984) for each locus as well as their variances. We estimated pairwise genetic differentiation between the *A. lacerata* sites by calculating *F_ST_* and *ρ_ST_* as proposed by Meirmans & Liu (2018) and Weir (1996). Finally, we used the Bayesian method implemented in ClonEstiMate (Becheler et al., 2017) and extended in GenAPoPop for polyploids (Stoeckel et al., 2024) to infer clonal rates using transition of genotype frequencies. We reported the 98% credible interval and the maximum posterior probability estimate from the distribution obtained for each population and successive temporal samples. We compared the change in genotype frequencies at the Maunalua Bay site between the two time points 2018 and 2019 in which we sampled blades. At ʻEwa Beach, Lagoon East, we inferred clonal rate using transition of genotype frequencies between samples from 2018-2019 and 2019-2021. As *Avrainvillea lacerata* is thought to be dioecious based on its phylogenetic position (Olsen-Stojkovich, 1985; Verbruggen et al., 2009), we did not infer a selfing rate because self-fertilization would not be possible if dioecious and may lead to a bias in clonal rates, especially if inbreeding occurred during sexual events.

Finally, we assessed genotypic richness (*R*) within each mound sampled as well as across all mounds in 2022 by determining the number of unique MLGs per mound. For the 2023 collection, we determined the number of MLGs found across two to three blades sampled per holdfast because some mounds visually appeared to have more than one holdfast.

## RESULTS

### Ploidy estimation

In an initial attempt to estimate ploidy, we used flow cytometry to evaluate DNA content of *Avrainvillea lacerata* nuclei. We employed a method of tissue dissociation and PI staining that successfully identified *Arabidopsis* nuclei for DNA content estimation but were unable to visualize *A. lacerata* nuclei using similar methods (Figure S1a-b). For *A. lacerata,* we obtained a single broad noise peak that we attributed to high levels of endogenous autofluorescence from photosynthetic pigments. With this high background fluorescence, we could not distinguish clear DNA positive peaks. We were, however, able to image *A. lacerata* DAPI-stained nuclei (Figure S2). The nuclei were in low abundance and in poor resolution, which prevented the analyses of DNA content. Moreover, we did not find gametes and would therefore have been unable to determine ploidy using this method.

### Microsatellite locus development

Of the 12 loci tested on the capillary sequencer, five loci produced peaks that were either too difficult to score confidently (e.g., stutters) or had poor amplification (Table S1). Seven loci amplified well by displaying clearly distinct alleles during initial simplex screening on the capillary sequencer. We amplified these seven loci in one multiplex and two duplexes (Table S2). Across the 321 genotyped blades, 175 blades (54.5%) displayed at least one locus with three or four different alleles (Table S3; Figure 2), 141 blades (43.9%) displayed a maximum of two alleles at all loci, and four blades (1.2%) had one allele at all seven loci. Overall, we concluded that the *Avrainvillea lacerata* blades we sampled were tetraploid (4N).

**Figure 2.**
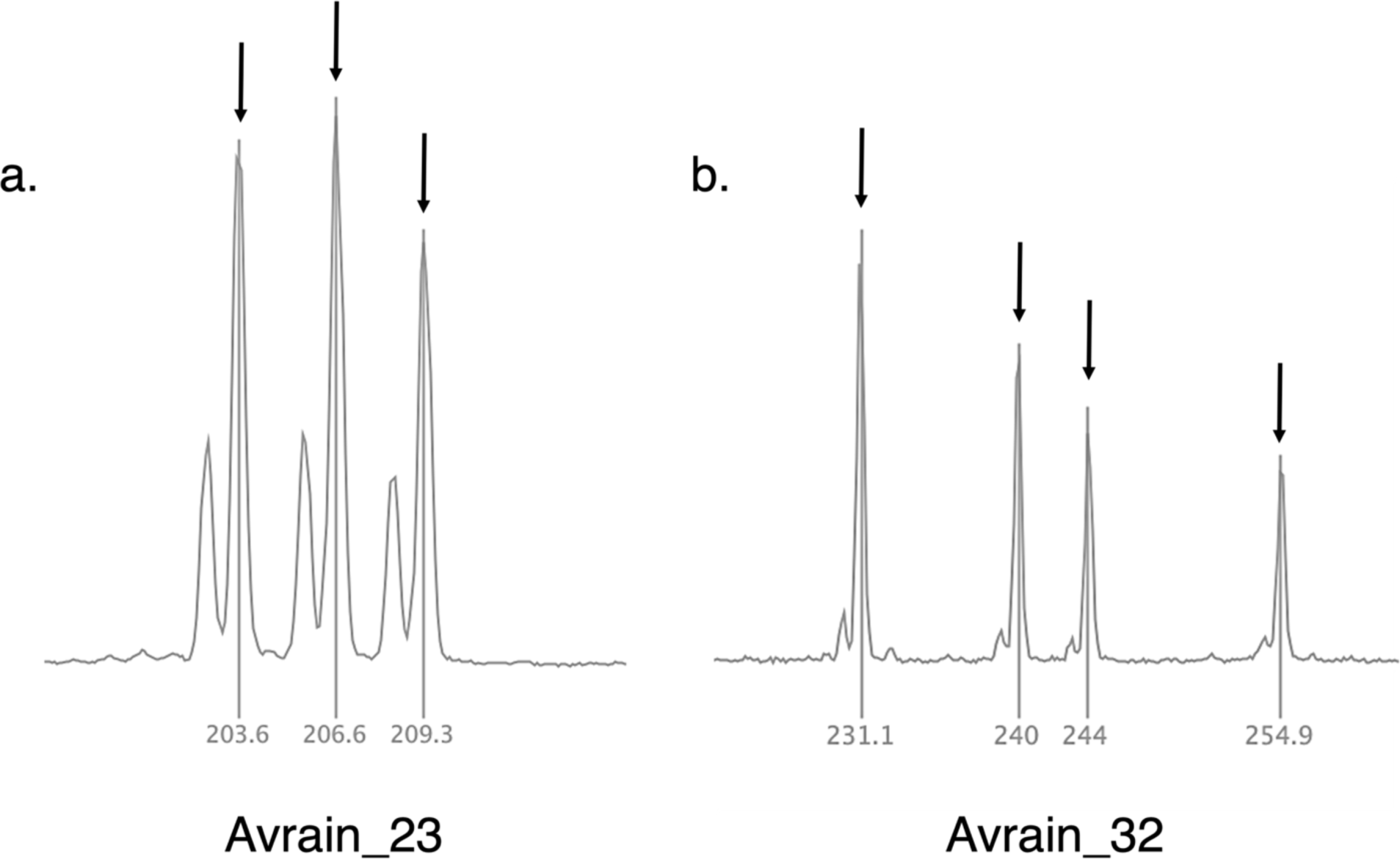
Fragment analysis traces for two microsatellite loci developed in *Avrainvillea lacerata*. Each arrow is pointing to a single allele. The architecture of these peaks are typical for microsatellites in which there is a short hitch before the larger peak. (a) Three different alleles are shown at locus Avrain_23. (b) Four different alleles are shown at locus Avrain_32.

### Allele dosage estimation

Blades that belonged to the same MLG based on raw alleles (i.e., the alleles found in the traces from fragment analysis) were assigned to new MLGs after allele dosage using the maximum likelihood method implemented in GenoDive (Table 1). In contrast, blades that belonged to the same MLG based on raw alleles more often were assigned to the same MLG when using allelic RFUs as a proxy for allele dosage (Table 1). Since estimating allele dosage using maximum likelihood artificially generated greater genotypic diversity, we conducted the population genetic analyses using genotypes generated by allelic RFUs.

**Table 1.** Number of MLGs per mound (N = 5 blades) and across all mounds (N = 40 blades total) sampled from ʻEwa Beach, Lagoon East during 2022 before and after allele dosage estimation using GenoDive and allelic relative fluorescence units (RFUs).

| Mound # | Number of MLGs based on raw alleles | Number of MLGs after allele dosage using maximum likelihood (Miermans, 2020) | Number of MLGs after allele dosage using RFU |
| --- | --- | --- | --- |
| 1 | 5 | 5 | 5 |
| 2 | 1 | 5 | 1 |
| 3 | 3 | 5 | 3 |
| 4 | 2 | 5 | 3 |
| 5 | 3 | 5 | 4 |
| 6 | 5 | 5 | 5 |
| 7 | 3 | 5 | 4 |
| 8 | 3 | 5 | 5 |
| All mounds | 25 | 40 | 30 |

### Genotypic and genetic diversity

At the Maunalua Bay site, in both 2018 and 2019, all blades sampled were unique MLGs (*R =* 1, *D\** = 1; Table 2). Observed heterozygosity (*H_O_*) varied between 0.405 and 0.643 in 2018 and between 0.473 and 0.755 in 2019 (Table 3). Expected heterozygosity (*H_E_*) varied between 0.499 and 0.749 in 2018 and between 0.480 and 0.750 in 2019 (Table 3). Single locus *F_IS_* values were all positive in 2018 and had greater variance as compared to 2019 where most values were closer to 0 (Table 3; Figure 3). *Pareto β* was > 2 in both 2018 and 2019 (Table 2). Linkage disequilibrium 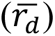 was 0.029 in 2018 and 0.004 during 2019 (Table 2). At Maunalua Bay, between 2018 and 2019, transition of genotypic frequency inferred rate of clonality (*c*) to fall within a credible interval superior to 98% that includes *c* = [0.41,0.68], with a maximum a posteriori probability estimated at *c* = 0.55 (Figure S3a).

**Figure 3.**
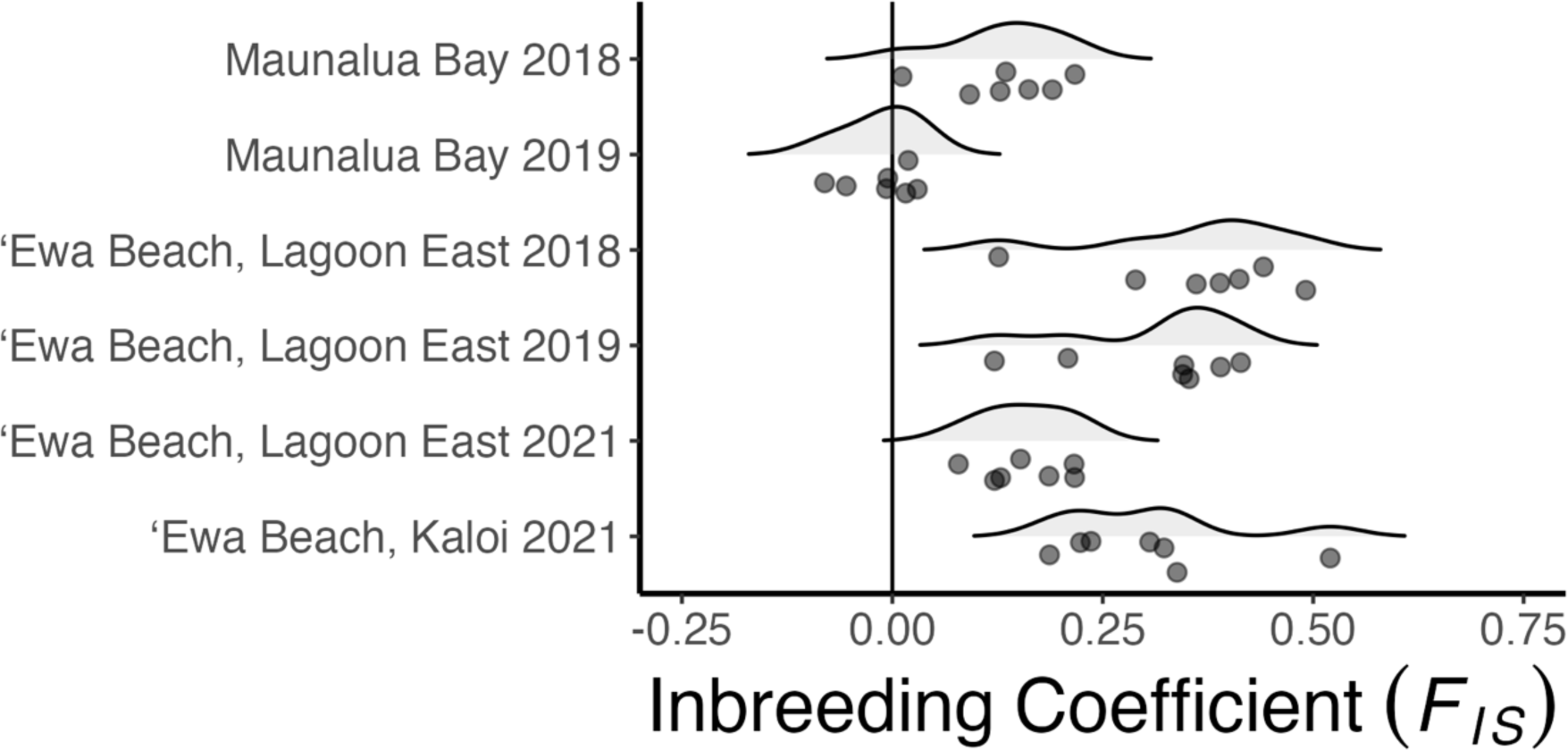
Density plots of single-locus values of the inbreeding coefficient (*F_IS_*) for each site from which *A. lacerata* was sampled in 2018, 2019, and 2021. The line at *F_IS_* = 0 shows Hardy-Weinberg proportions. Single locus *FIS* estimates are shown as points under the respective density plots (N = 7 loci).

**Table 2.** Genotypic richness (*R*), genotypic evenness (*D*\*), *pareto β*, and linkage disequilibrium 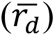 at the three sites in which we sampled *A. lacerata*.

|  | Maunalua Bay |  | ‘Ewa Beach, Lagoon East |  |  | ‘Ewa Beach, Kaloi |
| --- | --- | --- | --- | --- | --- | --- |
|  | 2018 | 2019 | 2018 | 2019 | 2021 | 2021 |
| Statistic | N = 49 | N = 49 | N = 47 | N = 49 | N = 30 | N = 30 |
| $R$ | 1.000 | 1.000 | 0.848 | 0.875 | 1.000 | 0.931 |
| $D^*$ | 1.000 | 1.000 | 0.985 | 0.982 | 1.000 | 0.995 |
| *Pareto* $\beta$ | 4.644 | 4.644 | 1.322 | 1.000 | 3.954 | 2.907 |
| $\overline{r_d}$ | 0.029 | 0.004 | 0.204 | 0.168 | 0.028 | 0.130 |

**Table 3.**
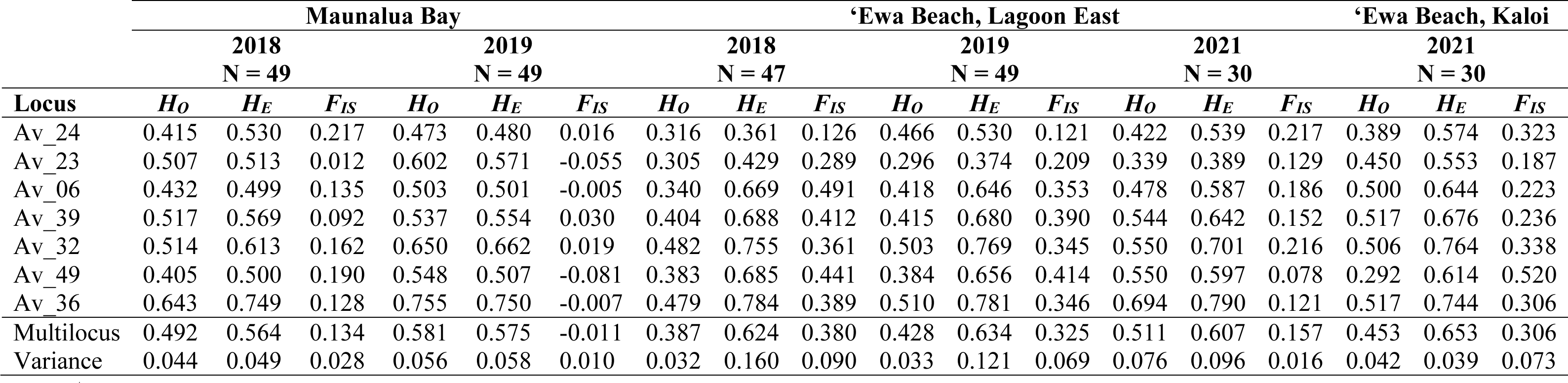
Single locus, multilocus, and variance for observed heterozygosity (*H_O_*), unbiased expected heterozygosity (*H_E_*), and the unbiased inbreeding coefficient (*F_IS_*) at the three sites in which we sampled *A. lacerata*.

At ʻEwa Beach, Lagoon East, in 2018, 2019, and 2021, genotypic richness (*R*) and evenness (*D\**) were high (*R* > 0.848; *D\** > 0.982), though both were less than 1 in 2018 and 2019 (Table 2). Observed heterozygosity (*H_O_*) varied between 0.305 and 0.482 in 2018, between 0.296 and 0.510 in 2019, and between 0.339 and 0.694 in 2021 (Table 3). Expected heterozygosity (*H_E_*) varied between 0.361 and 0.784 in 2018, between 0.374 and 0.781 in 2019, and between 0.389 and 0.790 in 2021 (Table 3). For all three time points, all single locus *F_IS_* were positive with the greatest variance in 2018 (Table 3; Figure 3). In 2018 and 2019, *pareto β* was between 0.7 and 2, respectively, but was > 2 in 2021 (Table 2). Lower levels of linkage disequilibrium 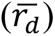 were observed at Lagoon East in 2021 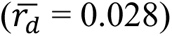 as compared to 2018 and 2019 (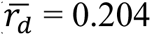 and 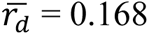, respectively) (Table 2). At Lagoon East, transition of genotypic frequencies inferred rate of clonality (*c*) between 2018 and 2019 to fall within a credible interval superior to 98% that includes *c* = [0.72, 0.86] with a maximum a posteriori probability estimated at *c* = 0.80, between 2019 and 2021 to fall within a credible interval superior to 98% that includes *c* = [0.45, 0.70] with a maximum a posteriori probability estimated at *c* = 0.58 (Figure S3b).

At ʻEwa Beach, Kaloi in 2021, both genotypic richness (*R*) and evenness (*D\**) were high (*R* = 0.931; *D\** = 0.995; Table 2). Observed heterozygosity (*H_O_*) varied between 0.292 and 0.517 (Table 3). Expected heterozygosity (*H_E_*) varied between 0.553 and 0.764 (Table 3). Single locus *F_IS_* were all positive and displayed greater variance than Lagoon East in 2021 (Table 3; Figure 3). *Pareto β* was > 2 (Table 2). Linkage disequilibrium 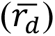 was 0.130 (Table 2).

### Genetic Differentiation

In 2018, pairwise *F_ST_* was 0.045 and *ρ_ST_* was 0.095 between Maunalua Bay and ʻEwa Beach, Lagoon East. In 2019, *F_ST_* was 0.025 and *ρ_ST_* was 0.065 between Maunalua Bay and ʻEwa Beach, Lagoon East. In 2021, *F_ST_* was 0.012 and *ρ_ST_* was 0.028 between ʻEwa Beach, Lagoon East and ʻEwa Beach, Kaloi.

### Within mound sampling

Across the eight mounds sampled in ʻEwa Beach, Lagoon East in 2022, we observed 30 MLGs out of 40 blades genotyped (Table 1). Mounds 1, 6, and 8 had blades that were all unique MLGs (Table 1). Mounds 3, 4, 5, and 7 had multiple MLGs, with some that we re-encountered more than once within the mound (Table 1). All blades from mound 2 belonged to the same MLG (Table 1).

Across all mounds sampled in 2023 and for which we determined the holdfast, we did not find any blades belonging to the same MLG (Figure S4). We observed multiple MLGs from the same putative holdfast (Figure S4). In some cases, MLGs were shared between blades sampled from different holdfasts within the same mound (e.g., Mound 2, MLG 235; Figure S4). In other cases, MLGs were shared between different mounds (e.g., Mound 2 & 3, MLG 234; Figure S4).

## DISCUSSION

We uncovered high clonal rates, between 55% to 80% depending on the years and sites, in *Avrainvillea lacerate,* supporting previous observations of vegetative mound growth (Peyton, 2009; Smith et al., 2002). This provides compelling genetic evidence that vegetative reproduction leads to the persistence and spread of *A. lacerata* at these two sites and likely elsewhere in the Hawaiian Archipelago. However, the *pareto β* values, low linkage disequilibrium, and positive *F_IS_* values hint at the occurrence of sexual reproduction, probably with significant inbreeding, despite gametes not being observed to date. When conducting within-mound analyses, we found evidence of what Olsen-Stojkovich (1979) referred to as ‘grafting’. Below, we discuss the ecological and evolutionary consequences of these findings and how they might contribute to the invasive potential of *A. lacerata* in Hawaiʻi.

### The reproductive system of *Avrainvillea lacerata*

Asexual reproduction has been assumed for other nuisance red algal species in the Hawaiian Archipelago, including *Acanthophora spicifera* (Smith et al., 2002)*, Gracilaria salicornia* (Smith et al., 2002; 2004)*, Hypnea musciformis* (Smith et al., 2002), and *Kappaphycus alvarezii* (Conklin & Smith, 2005; Smith et al., 2002), but has only been confirmed with population genetic data in *Chondria tumulosa* (Williams et al., 2024). Here, we show vegetative reproduction has played an important role in the spread of *Avrainvillea lacerata* in Hawaiʻi. Overall, our predictions for each of the two sites we sampled were supported, such that vegetative reproduction was greater at ‘Ewa Beach Lagoon East as compared to the site in Manaulua Bay. It is unclear whether clonal rates are greater in Hawaiʻi than in the native range (given that the native range is unknown) as has been documented in other macroalgae (e.g., *Caulerpa taxifolia,* Arnaud-Haond et al., 2017; *Gracilaria vermiculophylla,* Krueger-Hadfield et al., 2016). Future sampling throughout the extant range of *A. lacerata* is required to determine whether the prevailing reproductive mode has shifted following the invasion of the Hawaiian Archipelago.

Our population genetic data also hint that sexual reproduction is occurring. To our knowledge, no reproductive *A. lacerata* blades have been observed in Hawaiʻi. Sexual reproduction is thought to be holocarpic (Clifton, 1997, 2013; Clifton & Clifton, 1999; Hillis-Colinvaux, 1984; Olsen-Stojkovich, 1985), rendering observations challenging, particularly in an uncalcified alga where there is no remnant “skeleton” following gamete release. Furthermore, gamete production and release in holocarpic taxa can occur rapidly. For instance, in *Halimeda* spp., there is about a 36-hour timeframe from when a thallus becomes reproductive to when it releases gametes (Clifton & Clifton, 1999). If *A. lacerata* is holocarpic, reproduction could easily be missed if not observed during the narrow timeframe in which it likely occurs.

The positive single locus *F_IS_* values hint that inbreeding may occur when gametes are exchanged, particularly at the ʻEwa Beach sites. In the Bryopsidales, gametes are negatively buoyant (Clifton, 2013). Therefore, fertilization likely occurs close to the substrate or via settlement onto other nearby *Avrainvillea lacerata* mounds. This phenomenon likely favors mating among nearby *A. lacerata* blades limiting gene dispersal, such as observed in some corals (see as an example, Combosch & Vollmer, 2011). Inbreeding likely plays a larger role at the ʻEwa Beach sites than at Maunalua Bay. This may be explained by the regular manual removal of *A. lacerata* in Maunalua Bay by volunteer groups (Mālama Maunalua, https://www.malamamaunalua.org/). Thus, the distribution of *A. lacerata* clumps is smaller and patchier at the Maunalua Bay site than at the ʻEwa Beach sites, possibly providing less opportunity for inbreeding. Future population genetic studies on *A. lacerata* should include greater sampling density within and among sites as well as across the genome (e.g., single nucleotide polymorphisms, SNPs). These data should be paired with phenology studies to determine reproductive patterns (see as an example Krueger-Hadfield et al., 2023).

### The role of polyploidy influencing patterns of genetic and genotypic diversity

In a previous study examining the systematics of *Avrainvillea* in Hawaiʻi, there was some sequence differentiation (max 1.41% divergence) between *A. lacerata* blades that was correlated with the east (including Maunalua Bay) and west (including ʻEwa Beach) sides of Oʻahu based on *rbc*L and *tuf*A sequence data (Wade, 2019). These results suggest that two separate introductions of *A. lacerata* may have occurred in Hawaiʻi (Wade, 2019). Genetic differentiation as measured by allele frequencies (*F_ST_*) was greater between the Maunalua Bay site and ʻEwa Beach sites, corresponding with the east and west sides of the south shores of Oʻahu respectively, than between *A. lacerata* from the two sites at ʻEwa Beach. This also held true for *ρ_ST_*, which also measures genetic differentiation, but is independent of ploidy and the variable inheritance patterns across the loci of polyploid organisms (Meirmans & Liu, 2018; Ronfort et al., 1998). Therefore, repeated introductions might have played a role in establishing these patterns of genetic differentiation among sites in which *A. lacerata* is now found in Hawaiʻi. Alternatively, *Avrainvillea* is not thought to have long distance dispersal strategies (Littler & Littler, 1992), and there is likely a lack of gene flow between Maunalua Bay and ʻEwa Beach that further leads to genetic differentiation.

Genetic diversity (*H_E_*) was comparable between the site at Maunalua Bay and ʻEwa Beach, Lagoon East in 2018 and 2019. However, differences in *H_E_* could be minimal as estimates of *H_E_* are independent of the number of alleles and less sensitive to samples sizes from collection protocols or bottlenecks (Allendorf & Luikart, 2007). On the other hand, genotypic diversity is more strongly influenced by sample size and population bottlenecks, which could cause abrupt changes in allele frequencies (Allendorf & Luikart, 2007). This might be why differences in genotypic richness were more pronounced between the Maunalua Bay and ʻEwa Beach sites in which there was greater genotypic richness and *pareto ß* at Maunalua Bay. Greater genotypic richness at the Maunalua site perhaps is not surprising given that the clonal rate was lower as compared to ʻEwa Beach, Lagoon East. Additionally, it is possible that manual removal at Maunalua Bay provided space for new recruits to settle as adult thalli from nearby sites or zygotes produced from sexual reproduction.

Surprisingly, we found greater genotypic richness than expected if asexual reproduction dominates at a site. For example, in the red algal genus *Gracilaria* (reviewed in Krueger-Hadfield et al., 2021), genotypic richness was low in many sites dominated by asexual reproduction in the form of thallus fragmentation. We analyzed our data assuming *Avrainvillea lacerata* is tetraploid. However, it is possible that some blades could be diploid or triploid because 141 blades out of 321 (∼44%) had a maximum of two alleles and 125 blades (∼39%) had a maximum of three alleles across all loci, respectively. Yet, the likelihood of the distributions of allele frequencies at each site suggest at least tetraploidy. Ploidy levels should be assessed using complementary methods, such as microspectrophotometry or flow cytometry, as well as methods that would enable the determination of the number of copies of an allele (e.g., SNP genotyping).

We attempted to confirm ploidy levels in this study; however, we were limited in our ability to adapt a working protocol due to the lack of fresh blades which are preferred for DNA content analysis (Doležel et al., 2007). With the limited amount of fresh blades at our disposal, we were unable to isolate nuclei for flow cytometry using the buffers tested in this study. We had the most success in isolating nuclei and using fluorescence microscopy and DAPI using blades preserved in silica gel (Figure S2), though images were in poor resolution due to damage to the nuclei likely caused by silica gel preservation. Other studies have had similar difficulties with bryopsidalean algae (see Varela-Álvarez et al., 2012). With access to fresh blades, DAPI staining may be the best method for DNA content estimation in *Avrainvillea lacerata*. However, when nuclei are successfully isolated, a second problem arises as we do not have a reference of known ploidy, such as gametes. DAPI fluorescence intensity values must be compared to a reference such as this for accurate determination of ploidy (Doležel et al., 2007). Unfortunately, we are unable to do this with *A. lacerata* in Hawaiʻi as reproductive blades have yet to be documented (Smith et al., 2002).

It would not be surprising for *Avrainvillea lacerata* to be polyploid because polyploidy is common across the green algae, including within the Bryopsidales (Kapraun, 1994, 2005; Kapraun & Nguyen, 1994; Kapraun & Shipley, 1990; Varela-Álvarez et al., 2012), as well as some reds (e.g., *Porphyra,* Varela-Álvarez et al., 2018) and browns (e.g., *Fucus,* Preston et al., 2022). Polyploidy could be an important factor in the success of *A. lacerata* because it is often correlated with increased genotypic diversity (Soltis & Soltis, 1999; Parisod et al., 2010; te Beest et al., 2012). For example, in studies of mixed ploidy populations of angiosperms, tetraploid plants had increased levels of genotypic diversity relative to diploid plants of the same species (Baldwin & Husband, 2013; Brown & Young, 2000). This has been observed in macroalgae as well; high genotypic diversity was detected in invasive polyploid populations of *Asparagopsis taxiformis* (Andreakis et al., 2009). Polyploidy itself may incur greater genotypic diversity as the larger number of independent alleles at each locus allows for larger effective population sizes than a similar sized population of diploids (Luttikhuizen et al., 2007; Meirmans & Van Tienderen, 2013). Therefore, polyploids can harbor a greater amount of heterozygosity that is lost at a slower rate than in diploids (Moody et al., 1993). Additionally, increased mutation rates associated with polyploidization could further increase diversity (Otto, 2007). Therefore, high genotypic diversity is maintained, despite high clonal rates (e.g., Quarin et al., 2001). Moreover, additional genomic copies increase the masking of deleterious mutations (Comai, 2005; Parisod et al., 2010) and lower inbreeding depression relative to diploid populations, which could be beneficial when populations might be forced to undergo inbreeding after a bottleneck event, such as range expansion (Ronfort et al., 1998). Therefore, polyploids may possess increased genomic flexibility, allowing them to adapt and persist within new environments (Parisod et al., 2010).

### Origins of polyploidy in *Avrainvillea lacerata*

Polyploidy can arise through two processes. Allopolyploidy is the hybridization of two or more closely related species. Autopolyploidy results from the multiplication of the entire genome. Opportunities for hybridization are greater if multiple species within a genus co-occur in the same habitat, and this has been documented in fucoid seaweeds (see Sousa et al., 2019). Multiple species of *Avrainvillea* have been documented to co-occur (Atlantic: Littler & Littler, 1992; Littler et al., 2004; Indo-Pacific: Olsen-Stojkovich, 1979, 1985; Wai et al., 2009). If hybridization does occur between different species of *Avrainvillea,* it is most likely to occur within the native range, though genetic studies of species in the genus *Avrainvillea* are required to explore this idea. There are two species of *Avrainvillea* currently documented in Hawaiʻi: *A. lacerata* (Brostoff, 1989; Wade, 2019) and *A. erecta* (Wade et al., 2018). However, the two species are both morphologically and phylogenetically distinct (Wade et al., 2018). Therefore, it seems unlikely that hybridization between multiple species has occurred in Hawaiʻi. Alternatively, in *Caulerpa prolifera*, microsatellite genotyping has suggested that unreduced gametes have contributed to the polyploidy in the Mediterranean (Varela-Álvarez et al., 2012). If gametes can be isolated from *A. lacerata* in Hawaiʻi, then we can determine the ploidy of the gametes and either support or refute the possibility of autopolyploidy in *A. lacerata*.

### *Avrainvillea lacerata* ‘individuals’

Coalescence, or the fusion between two or more individuals generating one entity composed of multiple genotypes (e.g., a chimera), has mostly been described in red algae, where some authors have suggested it is widespread and occurs naturally *in situ* (Santelices et al., 1999; Santelices et al., 2003). It seems to incur numerous ecological benefits, such as increased survival (Santelices, et al., 2003; Santelices et al., 1999). Coalescence was described in another bryopsidalean green alga, *Codium* sp., when González & Santelices (2008) found that separate, but neighboring crusts often fused together. This was supported after screening thalli with a chloroplast gene (Trn-Gly) when single crusts were shown to include multiple haplotypes (González & Santelices, 2008).

Olsen-Stojkovich (1979) described ‘grafting’ in *Avrainvillea* spp. in which juveniles may grow fused together. This is perhaps not surprising given the lateral spread of subterranean holdfasts (Littler et al., 2004, 2005; Peyton, 2009). Grafting is a term often used in horticultural practices to describe when two different parts of a plant are joined together to fuse and continue their growth in such a way that intercellular connections form (Melnyk & Meyerowitz, 2015). These techniques were inspired by the processes of natural grafting when stems or roots of plants attach and fuse together (Mudge et al., 2009). Grafting is used to adjust the size of plants, improve stress tolerance, or allow plants to grow in different environments (Melnyk & Meyerowitz, 2015). This definition of grafting differs from the coalescence described in *Codium* and other red algae. Rather, coalesced filaments of separate *Codium* crusts grew intermixed without establishing intercellular connections with one another (González & Santelices, 2008). Likewise, *Chondrus crispus* forms large holdfasts that can have female gametophytes, male gametophytes, and tetrasporophytes, but to date no chimeric fronds have been genotyped (see Krueger-Hadfield, 2011; Krueger-Hadfield et al., 2013). Instead, holdfasts may coalesce, enhancing survival, without exchanging genetic material (Krueger-Hadfield, 2011). Currently, it is unclear whether fusions between *Avrainvillea* thalli are composed of intercellular connections or intermixed filaments, but we cannot discount the possibility of either.

We found multiple genotypes within single mounds, including prior to estimating allele dosage (i.e., from raw alleles found in fragment analysis traces), in *Avrainvillea lacerata*. These results suggest *A. lacerata* might be a mosaic structural unit (see discussion in Clarke, 2012). Moreover, we found distinct genotypes when blades clearly shared the same holdfast. Until we have an independent measure of ploidy, we cannot fully discount the possibility that some blades are chimeras following the ‘grafting’ described by Olsen-Stojkovich (1979) and the subsequent coalescence of siphons. However, the distribution of allelic RFUs and allele frequencies corresponded to the multinomial sampling of a standard tetraploid. This would have not been the case in chimeras that would have shown all the possible distributions of RFUs depending on the relative importance of each different genetic line. This suggests that most thalli are at least tetraploid. Future studies need to use field experimentation, population genetics, and histological studies to confirm the coalescence of siphons in *Avrainvillea*.

Alternatively, genotypic variation within a mound does not necessarily mean that the mound is composed of multiple genets or “individuals”. One major misconception in distinguishing individuals lies in the assumption that all parts that make up the individual genet, including ramets and modules of modular organisms, are genetically homogenous (Scrosati, 2002). In clonal organisms that undergo indeterminate growth and can have relatively long lifespans, such as macroalgae, somatic mutations occur during vegetative growth resulting in a mosaic of genotypes within a single genet (Gill et al., 1995). This may be especially relevant in microsatellites, which generally have higher mutation rates (Anmarkrud et al., 2008). Ramets produced from the same genet could display genetic variation, generating multiple levels in which natural selection could potentially act upon (Clarke, 2012; Tuomi & Vuorisalo, 1989). If we define a genet as a distinct MLG, it is possible that we might erroneously assume two *Avrainvillea lacerata* blades with different MLGs as being from separate genets. Instead, it is possible they could have come from the same zygote and somatic mutations have generated a genetic mosaic over time. Thus, it is possible that the genotypic heterogeneity within *Avrainvillea* mounds might not be the result of coalescence of different ‘individuals’ but rather an accumulation of somatic mutations within a single genet during vegetative growth of a mound.

### Past removal efforts and critical next steps for the management of *Avrainvillea lacerata*

Due to its rapid spread (Smith et al., 2002; Veazey et al., 2019; Wade, 2019) and negative effects on native seagrass beds (Peyton, 2009) and intertidal benches (Foster et al., 2019), finding a way to effectively combat the spread of *Avrainvillea lacerata* is a high priority for local management agencies. Understanding the basic biology of *A. lacerata* is essential with which to develop management strategies, including an understanding of the spread of this alga (Zamora et al., 1989). *A. lacerata* primarily spreads via vegetative growth of mounds, which suggests the total removal of biomass would be critical for species management. Non-profit organizations, such as Mālama Maunalua, continue to recruit volunteers for manual removal of *A. lacerata* in Maunalua Bay (https://www.malamamaunalua.org/). In 2010, the U.S. government provided a large award that allowed a massive restoration project at Maunalua Bay (Kittinger et al., 2016). During this removal effort, 1.32 million kg of non-native macroalgae, including *A. lacerata,* were removed by hand and transported to a composting site (Kittinger et al., 2016). This removal effort did appear to have some short-term success as the community composition of cleared areas seemed to resemble those of native plots (Longenecker et al., 2011). Large-scale manual removal efforts can substantially reduce the standing biomass, but holdfasts left behind can result in regrowth via vegetative spread of *A. lacerata* mounds. If manual removal is used, any fragments left behind need to be small enough to hinder lateral spread of holdfasts and blade production (see discussion in Bonnett et al., 2014). Smith et al. (2002) showed that fragments of *A. lacerata* holdfasts that were at least three centimeters showed the highest success for regrowth. Therefore, future experiments should examine whether fragments smaller than three centimeters are able to regrow in the field in order to confirm the maximum fragment size that would hinder regrowth of *A. lacerata*. This is especially critical as it is likely impossible to remove all fragments of *A. lacerata*. Our data from Manaulua Bay suggest the regrowth of some genotypes that were left behind as well as the potential of new recruitment from juveniles or adults.

Large-scale chemical treatment could potentially overcome the issue of leaving behind thalli by causing mortality of entire *Avrainvillea lacerata* mounds. Van De Verg and Smith (2022) showed that injecting *A. lacerata* mounds with small amounts hydrogen peroxide decreased photosynthetic efficiency, potentially leading to a decrease in biomass. While the immediate effects of injection seem promising, the long-term effects of this form of control remain to be tested and large-scale application appears impractical. For instance, chemical treatments aiming to control invasive *Caulerpa taxifolia* have been shown to be harmful to animal life within the treatment area (Williams & Schroeder, 2004). This is particularly pertinent given the faunal community found within *A. lacerata* holdfasts, which is composed of native species (Magalhães & Bailey-Brock, 2014). It is also unclear whether this treatment can be applied to other sites where *A. lacerata* is present. Experimentation of this method took place in the more protected, soft sediment habitat of Maunalua Bay (Van De Verg & Smith, 2022), but the logistics of effectively applying this across the diverse habitats *A. lacerata* inhabits, including mesophotic regions (Spalding et al., 2019) and more wave exposed rocky shorelines (Foster et al., 2019) where *A. lacerata* is found, will likely be limited.

Prior to this study, sexual reproduction in *Avrainvillea lacerata* in Hawaiʻi was not considered for population growth based on anecdotal absence of observable gametangia. Additional research is needed to understand sexual reproduction in this alga. For example, phenology studies are required to examine when and how often sexual reproduction occurs and the environmental triggers for gametogenesis. Phenology studies of bryopsidalean macroalgae have occurred in Caribbean coral reefs (Clifton & Clifton, 1999), but these studies did not include members of the genus *Avrainvillea.* However, for the genera that were studied (i.e., *Caulerpa, Halimeda, Penicillus, Rhipocephalus,* and *Udotea*) several instances of sexual reproduction occurred throughout the seasonal peak for reproductive activity (Clifton & Clifton, 1999). Light levels and water temperature seemed to have some influence on the timing of reproductive activity as the seasonal peak of reproduction corresponded with the shift from the dry season to the wet season, a period of increased solar radiation (Clifton & Clifton, 1999). If the timeframe and environmental conditions of sexual reproduction for *A. lacerata* are determined, then biomass removal efforts could be prioritized before thalli become reproductive. This is similar to the logic behind suggestions for management in controlling the spread of invasive angiosperms where sexual reproduction and seed dispersal are considered to be a mechanism for spread (Bonnett et al., 2014). The dispersal of sexual propagules, including gametes and zygotes, could be one of the mechanisms that has allowed *A. lacerata* to spread across the southern shores of Oʻahu. If this is the case, then sexual reproduction at nearby sites could result in the dispersal and recruitment of new individuals into sites that have been cleared.

Understanding the dispersal of *Avrainvillea lacerata* in Hawaiʻi is is crucial to minimizing or preventing future movement of invasive species (Wilson et al., 2009). If *A. lacerata* can disperse across localities, removal efforts will need to expand beyond Maunalua Bay. By understanding the connectivity of *A. lacerata* at sites across the Hawaiian Archipelago, we can aid in the development of better management strategies. For example, angiosperms with long distance dispersal mechanisms (i.e., wind-based dispersal) display less genetic differentiation, and therefore greater connectivity, among sites than plants with shorter distance dispersal (Govindaraju, 1988). We assessed connectivity between Maunalua Bay and ʻEwa Beach by estimating pairwise genetic differentiation based on allele identity (*F_ST_*) and allele size (*ρ_ST_*). Based on both measurements of genetic differentiation, the level of divergence was greater between Maunalua Bay and ʻEwa Beach, Lagoon East than it was between the two sites at ʻEwa Beach. This suggests some isolation by distance, but more sites should be sampled to sufficiently test for a correlation between genetic and geographical distances. It is important to emphasize that this differentiation may also be due to separate colonization events, if *Avrainvillea lacerata* is indeed non-native to Hawai‘i. The connectivity of *A. lacerata* among sites in Hawaiʻi should be explored further by including broader sampling along the shores of Oʻahu, at the neighboring islands of Maui and Kauai, and across its broad depth gradient (intertidal to mesophotic depths) Then, the combined understanding of reproduction and connectivity can aid managers in developing the most effective management plans for a given site and more widely throughout the Hawaiian Archipelago.

Finally, the origin of *Avrainvillea lacerata* in Hawaiʻi is unresolved. The alga may have been introduced from Japan based on the close phylogenetic grouping of *A. lacerata* from Hawaiʻi with specimens from Japan (Wade, 2019). Alternatively, due to its high abundance in the mesophotic as well as the absence of mesophotic studies prior to the initial collection of *A. lacerata*, it is possible that *A. lacerata* is native in Hawaiʻi and existed in the mesophotic prior to invading the intertidal (Spalding, 2012). It is also possible that *A. lacerata* was introduced from Guam. A different Hawaiian invader, *Acanthophora spicifera,* was introduced to Hawaiʻi through increased shipping traffic between Honolulu and Guam during World War II (Doty, 1961). It is plausible that *A. lacerata* could have been introduced in a similar manner, and perhaps invaded the mesophotic prior to invading the intertidal in the 1980’s. Future work needs to include sampling *A. lacerata* broadly across the Indo-Pacific and the Hawaiian Archipelago to determine the source populations of *A. lacerata* in Hawaiʻi (see, as example, Krueger-Hadfield et al., 2017). If the source of the *A. lacerata* invasion can be determined, comparisons can then be made to understand how the niche of *A. lacerata* may have shifted in the invasion of the Hawaiian Islands. Invasion studies often compare the non-native range to the entirety of the native range (see Petitpierre et al., 2012). However, not all native sites contribute equally to an invasion, thereby leading to an underestimation of adaptation, inaccurate inferences of the direction of evolutionary change, or both. For example, when comparing the niche of non-native *Gracilaria vermiculophylla* thalli to the entirety of the native range, it seemed that non-native thalli experienced niche expansion into colder temperatures (Sotka et al., 2018). However, when compared to the source populations, non-native thalli expanded into warmer habitats (Sotka et al., 2018). Therefore, identifying the source populations will improve the ability to detect the evolutionary shifts that may have allowed *A. lacerata* to proliferate in Hawaiʻi.

### Conclusions and future directions

The microsatellite data from this study highlight the importance of the most basic biological information, such as the ploidy level, to confidently analyze population genetic data. Analyzing DNA content in relation to a reference of known ploidy will be a critical next step for further exploration of the population genetic structure of *Avrainvillea lacerata* in Hawaiʻi and throughout its extant range. Recent advances in genotyping methods that exploit increased sequencing depth with low error rates have unlocked this possibility for population genetic analyses of polyploid species (e.g., Hi-Plex genotyping with single nucleotide polymorphisms (SNPs); Delord et al., 2018) and may be an avenue to pursue to confidently assign allele dosage (see Stoeckel et al., 2024).

The rate of clonality in *Avrainvillea lacerata* in Hawaiʻi is high. Asexual reproduction plays an important role in the spread of many nuisance macroalgae in the Hawaiian Archipelago (Conklin & Smith, 2005; Smith et al., 2002, 2004; Williams et al., 2024). Our genetic data and previous ecological obersvations at Maunalua Bay suggest that removal efforts may not remove all of the holdfast material, although the downstream consequences of this are unclear. The microsatellite loci used in this study will facilitate future studies that are necessary to understand the invasive history of this alga. More broadly, this work adds to the body of literature for green algae in which we know very little regarding the population genetic patterns (Krueger-Hadfield et al., 2021), despite the work of Otto and Marks (1996) highlighting the role green algae can play in unlocking the secrets of eukaryotic sex.

## ACKNOWLEDGEMENTS

We thank Our Project in Hawaiʻi’s Intertidal (OPIHI), Mālama Maunalua, L. Morrow, J. Dagostino, N. Strait, H. Fullerton, G. Kuba, S. Bruna, C. Connolly, B. Connolly, and T. Williams for field help; Haseko Development, Inc. for their support of algal research in ʻEwa as part of their intertidal sampling program; J. Aida for initial help with microsatellite screening; C. Amsler for serving on BMT’s MS thesis committee and providing feedback on this manuscript; A. Sherwood for providing initial material for microsatellite development; K. Mukhtar and R. Bedgood for providing *Arabidopsis* for flow cytometry; M. Crowley, M. Han, and A. Cao for help with fragment analysis at the University of Alabama at Birmingham (UAB) Heflin Center for Genomic Services; and B. Kennedy for access to the Bernice P. Bishop Museum Herbarium Pacificum and library. This project was supported by Alabama Academy of Science student research award (to BMT), the Phycological Society of America Grant In Aid of Research (to BMT), the UAB Graduate Student Government (GSG) Professional Development and Travel Award (to BMT), the UAB Harold Martin Outstanding Student Development Award (to BMT), CLONIX-2D (ANR-18-CE32-0001 to SS and SAKH), start-up funds from the College of Arts and Sciences at UAB (to SAKH and MLH), the National Science Foundation (NSF) DEB-2113745 (to SAKH), and National Fish and Wildlife Foundation (#74235 to SAKH and HLS). SAKH was supported by the NSF CAREER award DEB-2141971. This project represents the partial fulfillment of the Master of Science degree for BMT.

## DATA ACCESSIBILITY

Genetic data and code are provided on Zenodo (LINK HERE WHEN DONE)

## AUTHOR CONTRIBUTIONS

**Brinkley M. Thornton** Data curation (equal), Formal analysis (equal), Funding acquisition (equal), Investigation (equal), Methodology (equal), Visualization (equal), Writing – original draft (equal), Writing – review and editing (equal); **Heather L. Spalding** Conceptualization (equal), Funding acquisition (equal), Investigation (equal), Resources (equal), Supervision (equal), Writing – review and editing (equal); **Solenn Stoeckel** Formal analysis (equal), Funding acquisition (supporting), Investigation (equal), Methodology (equal), Resources (equal), Software (lead), Writing – review and editing (equal); **Melissa L. Harris** Investigation (equal), Methodology (equal), Resources (equal), Supervision (equal), Writing – review and editing (equal); **Rachael M. Wade** Investigation (supporting), Writing – review and editing (equal); **Stacy A. Krueger-Hadfield** Conceptualization (equal), Data curation (equal), Funding acquisition (lead), Investigation (equal), Methodology (equal), Project Administration (lead), Resources (lead), Software (supporting), Supervision (lead), Visualization (equal), Writing – original draft (equal), Writing – review and editing (equal)

## ABBREVIATIONS

MLG: Multilocus Genotype
PCR: Polymerase Chain Reaction
RFU: Relative Fluorescence Units

## SUPPLEMENTAL TABLES

**Table S1.**
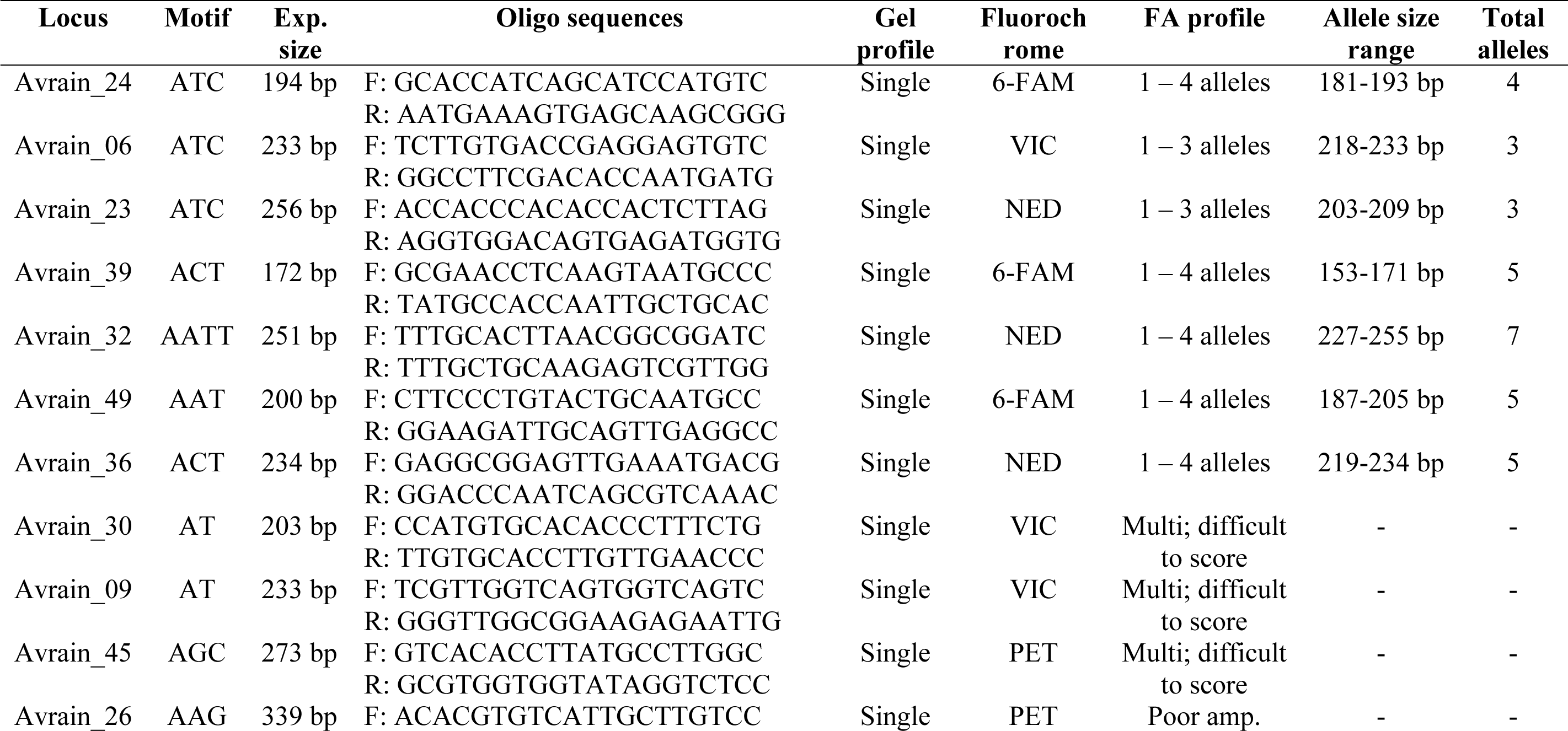
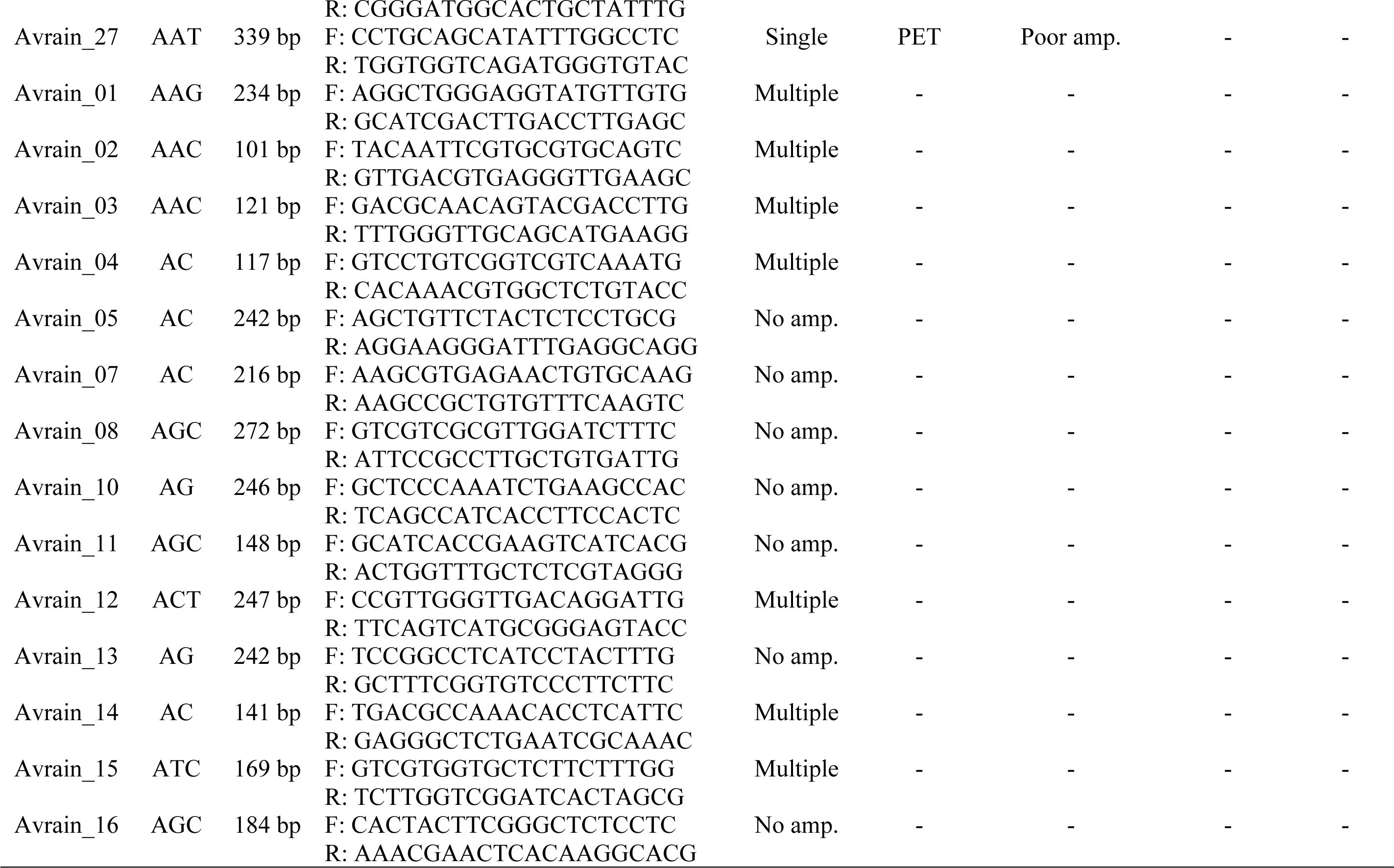
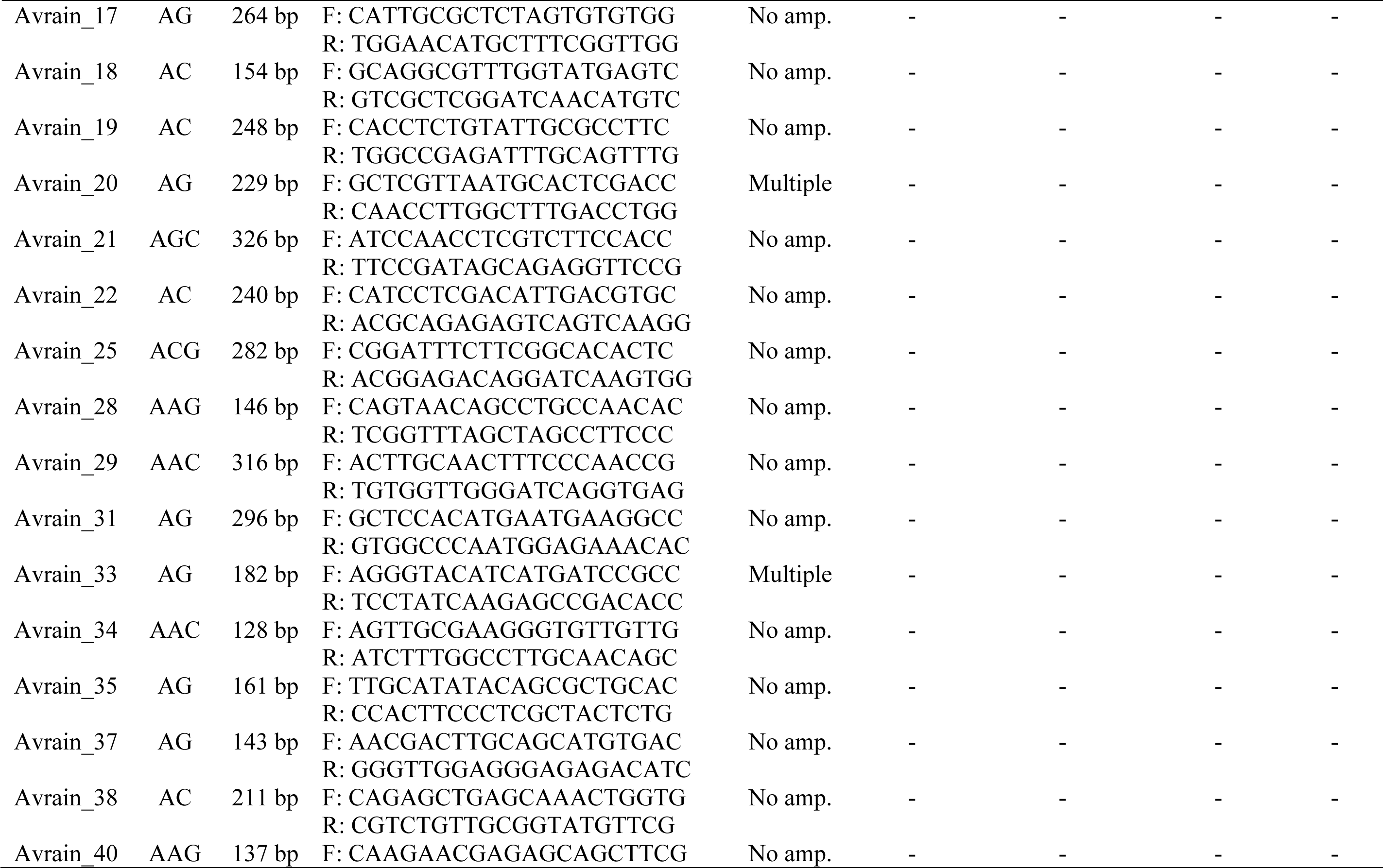
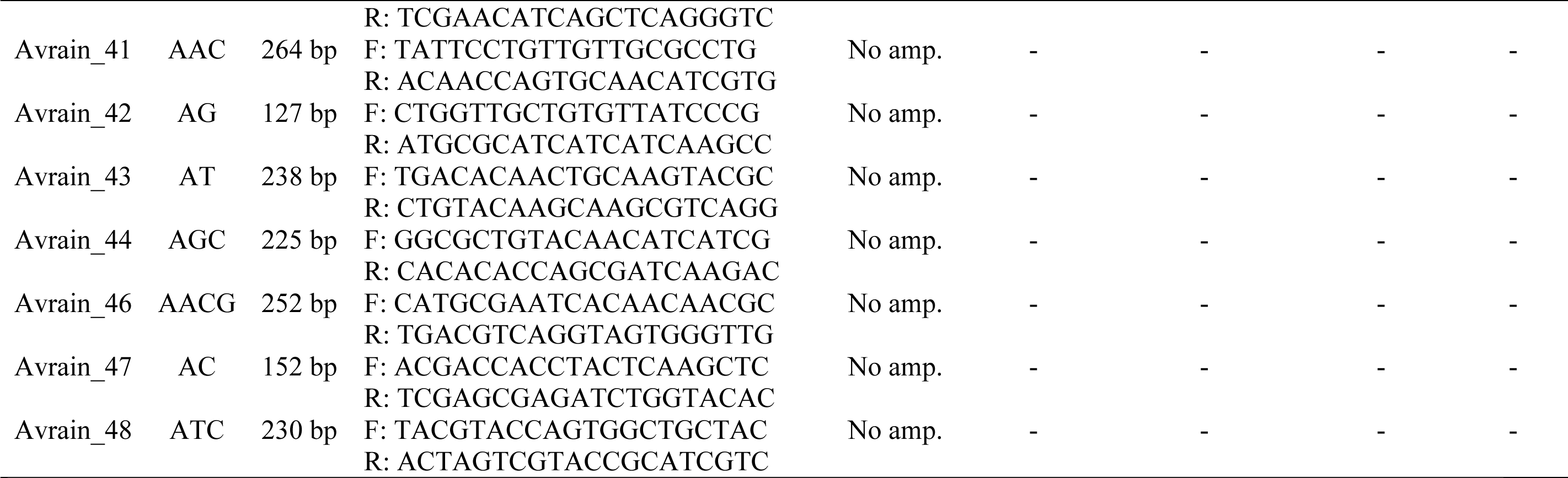
Microsatellite locus information for *Avrainvillea lacerata* in Hawaiʻi. Locus name, repeat motif, expected size, forward and reverse oligo sequences, agarose gel amplification profile, forward oligo fluorochrome, and fragment analysis (FA) amplification profile. Note fluorochrome and FA profile columns are only for loci tested on the capillary sequencer.

**Table S2.** *Avrainvillea lacerata* microsatellite loci. Multiplex pri mer concentrations and fluorochrome.

| Locus | Multi- or duplex | Dye | Concentration of<br>labeled forward oligo | Concentration of<br>unlabeled forward oligo | Concentration of<br>reverse oligo |
| --- | --- | --- | --- | --- | --- |
| Avrain_24 | M1 | 6-FAM | 250 nM | 200 nM | 450 nM |
| Avrain_06 | M1 | VIC | 200 nM | 100 nM | 300 nM |
| Avrain_23 | M1 | NED | 225 nM | 125 nM | 350 nM |
| Avrain_39 | D1 | 6-FAM | 200 nM | 100 nM | 200 nM |
| Avrain_32 | D1 | NED | 225 nM | 125 nM | 350 nM |
| Avrain_49 | D2 | 6-FAM | 225 nM | 125 nM | 350 nM |
| Avrain_36 | D2 | NED | 225 nM | 125 nM | 350 nM |

**Table S3.** The proportion of more than 2 alleles at *Avrainvillea lacerata* microsatellite loci across all thalli genotyped (N=321)

| Locus | Frequency of thalli with 3 alleles | Frequency of thalli with 4 alleles |
| --- | --- | --- |
| Avrain_24 | 39 | 2 |
| Avrain_06 | 13 | 0 |
| Avrain_23 | 78 | 0 |
| Avrain_39 | 48 | 1 |
| Avrain_32 | 43 | 18 |
| Avrain_49 | 22 | 0 |
| Avrain_36 | 70 | 38 |

## SUPPLEMENTAL FIGURES

**Figure S1.**
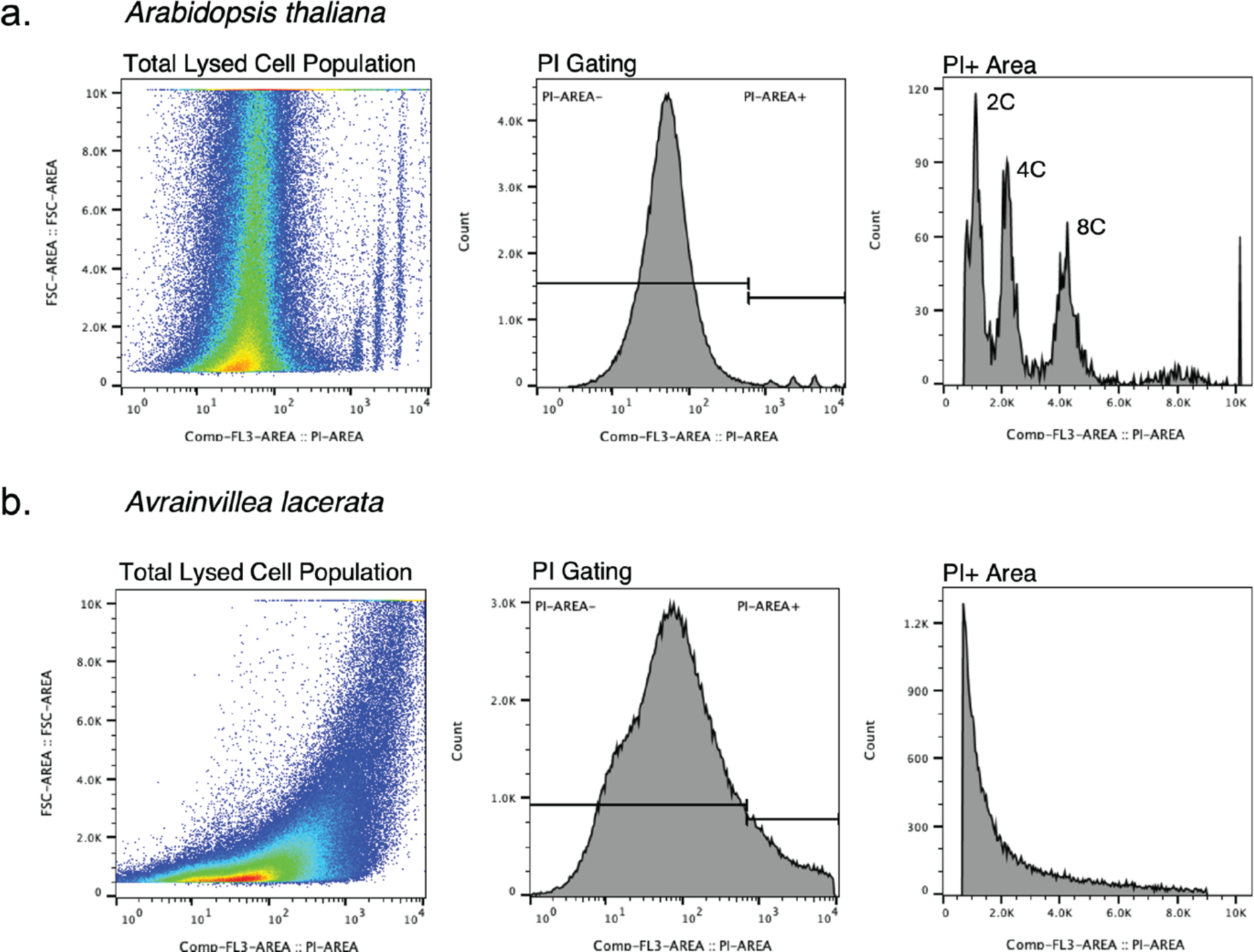
Flow cytometry data for *Arabidopsis thaliana* (a) and *Avrainvillea lacerata* (b) lysed using LB01. (a) Left graphs: forward scatter vs propidium iodide (PI) area in log form for the total dissociated tissue population. Center graphs: histograms demonstrating gating strategy for PI-negative (PI-AREA-) and PI-positive (PI-AREA+) staining on log scale (center graphs). Right graphs: histograms of PI-positive staining on linear scale demonstrating the direct relationship between fluorescence intensity and DNA content. Three prominent PI+ DNA peaks are observed in *Arabidopsis thaliana* cell lysis preparations. No distinct PI+ peaks are visible in *Avrainvillea lacerate* cell lysis preparations. Peaks represent nuclei in di fferent stages of the cell cycle. 2C=G_1_; 4C=G_2_; 8C=somatic polyploidy.

**Figure S2.**
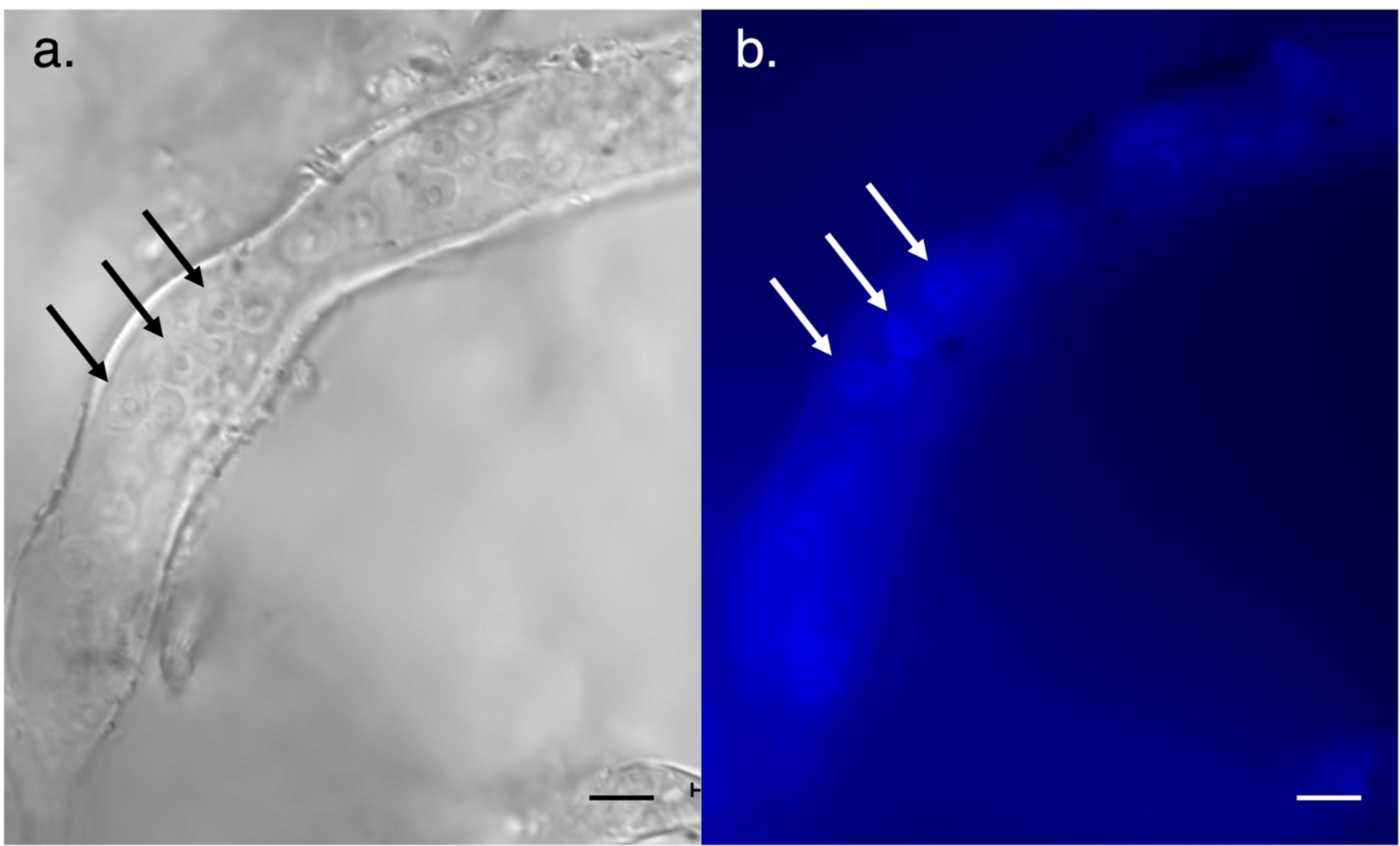
(a) Brightfield image of *A. lacerata* siphon where nuclei were isolated. Scalebar: 100 μm. (b) Fluorescent imaging of same *A. lacerata* siphon showing DAPI stained nuclei (region circled in white). Nuclei are indicated by oval to circular shaped regions of homogenous DAPI fluorescence with some nuclei showing an apparent nucleolus. Scalebar: 100 μm.

**Figure S3.**
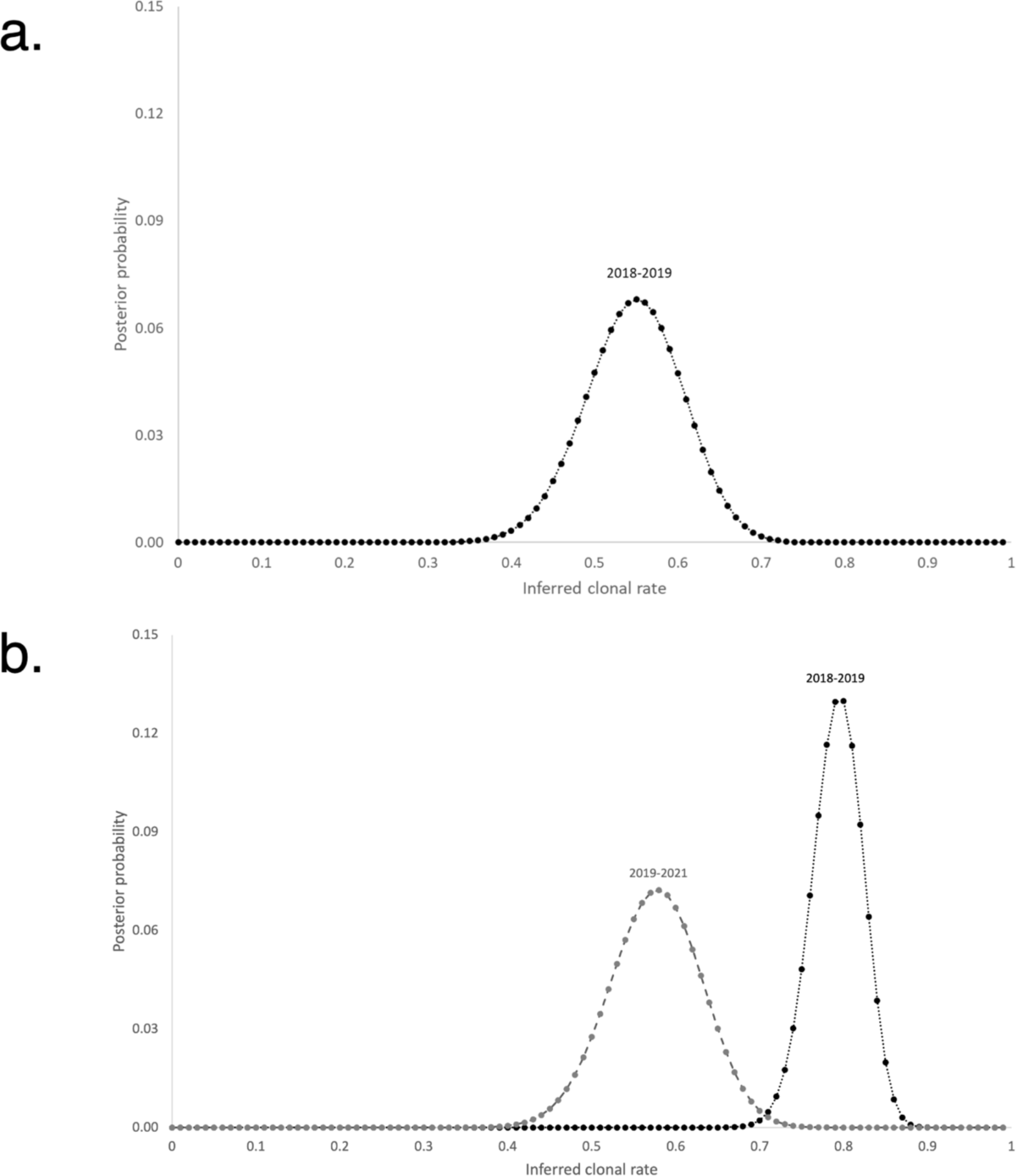
Posterior probabilities of rates of clonality using changes of genotype frequencies between temporal samples (CEMP). (a) CEMP Plot for Maunalua Bay between 2018 and 2019. (b) CEMP Plot for ʻEwa Beach, Lagoon East between 2018 and 2019 (black posterior distribution, dotted line) and 2019-2021 (dark grey posterior distribution, dashed line).

**Figure S4.**
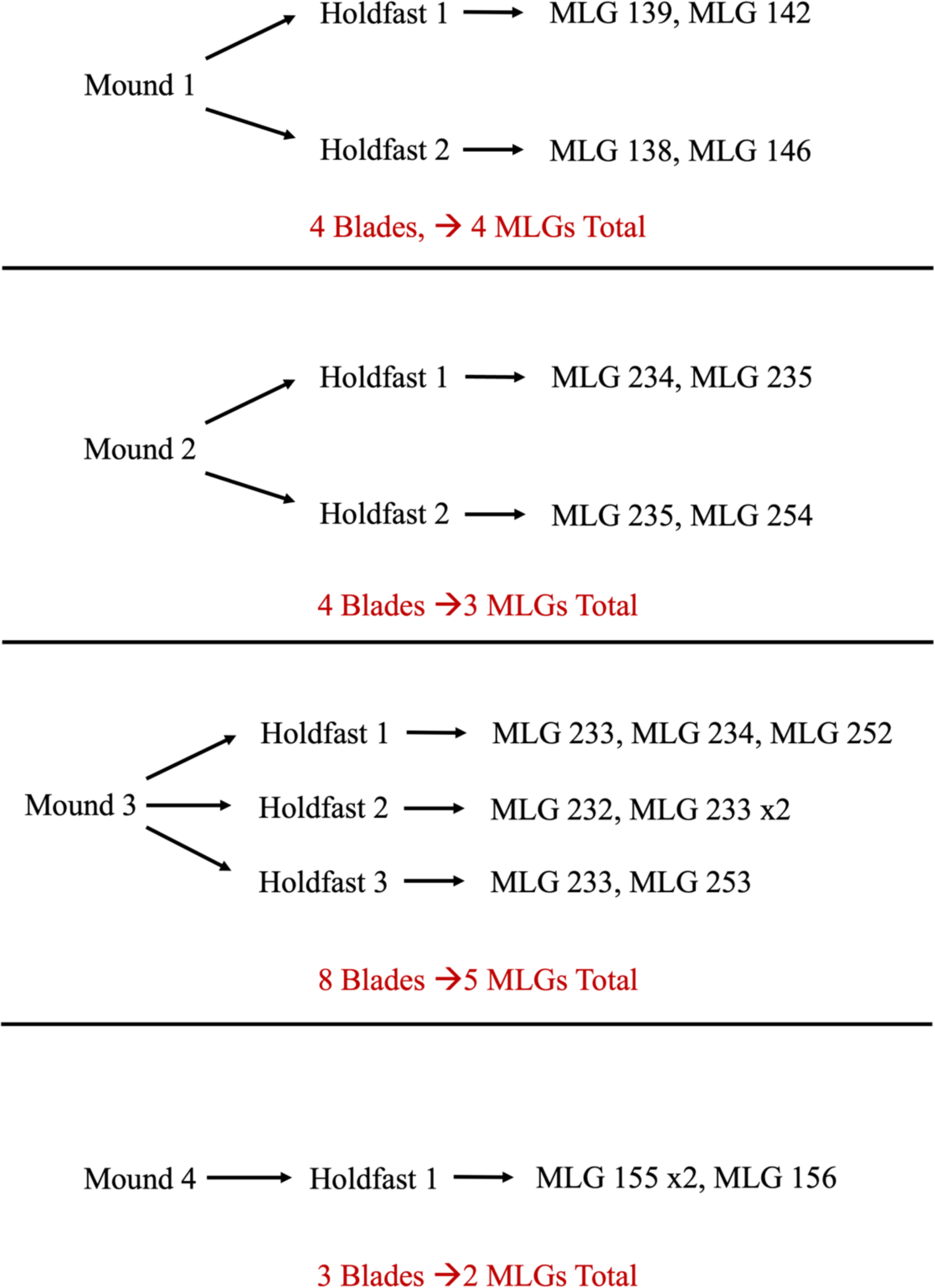
Breakdown of MLGs found in mounds sampled from ʻEwa Beach, Lagoon East during 2023. MLG numbers were assigned to individual blades based on their MLG by GenAPoPop (Stoeckel et al., 2024).

## REFERENCES

Agapow, P., & Burt, A. (2001). Indices of multilocus linkage disequilibrium. Molecular Ecology Notes, 1, 101–102.

Allendorf, F. W., & Luikart, G. (2007). Conservation and the genetics of populations. Blackwell Publishing Ltd.

Andreakis, N., Kooistra, W. H. C. F., & Procaccini, G. (2009). High genetic diversity and connectivity in the polyploid invasive seaweed *Asparagopsis taxiformis* (Bonnemaisoniales) in the Mediterranean, explored with microsatellite alleles and multilocus genotypes. Molecular Ecology, 18, 212–226.

Anmarkrud, J. A., Kleven, O., Bachmann, L., & Lifjeld, J. T. (2008). Microsatellite evolution: Mutations, sequence variation, and homoplasy in the hypervariable avian microsatellite locus HrU10. BMC Evolutionary Biology, 8, 138.

Antonovics, J. (1968). Evolution in closely adjacent plant populations: V Evolution of self-fertility. Heredity, 23, 219–238.

Arnaud-Haond, S., Aires, T., Candeias, R., Teixeira, S. J. L., Duarte, C. M., Valero, M., & Serrão, E. A. (2017). Entangled fates of holobiont genomes during invasion: Nested bacterial and host diversities in *Caulerpa taxifolia*. Molecular Ecology, 26, 2379–2391.

Arnaud-Haond, S., Duarte, C. M., Alberto, F., & Serrão, E. A. (2007). Standardizing methods to address clonality in population studies. Molecular Ecology, 16, 5115–5139.

Baker, H. G. (1955). Self-compatibility and establishment after “long-distance” dispersal. Evolution, 9, 347–349.

Baldwin, S. J., & Husband, B. C. (2013). The association between polyploidy and clonal reproduction in diploid and tetraploid *Chamerion angustifolium*. Molecular Ecology, 22, 1806–1819.

Barrett, S. C. H. (2008). Major evolutionary transitions in flowering plant reproduction: An overview. International Journal of Plant Sciences, 169, 1–5.

Barrett, S. C. H. (2011). Why reproductive systems matter for the invasion biology of plants. In D. M. Richardson (Ed.), Fifty Years of Invasion Ecology (1st ed., pp. 195–210). Blackwell Publishing Ltd.

Becheler, R., Masson, J., Arnaud-Haond, S., Halkett, F., Mariette, S., Guillemin, M., Valero, M., Destombe, C., & Stoeckel, S. (2017). ClonEstiMate, a Bayesian method for quantifying rates of clonality of populations genotyped at two-time steps. Molecular Ecology Resources, 17, e251–e267.

Bell, G. (1994). The comparative biology of the alternation of generations. In M. Kirpatrick (Ed.), Lectures on mathematics in life sciences: The evolution of haplo-diploid life cycles (pp. 1–26). American Mathematical Society.

Bellard, C., Cassey, P., & Blackburn, T. M. (2016). Alien species as a driver of recent extinctions. Biology Letters, 12, 20150623.

Bierzychudek, P. (1985). Patterns in plant parthenogenesis. Experientia, 41, 1255–1264.

Bonnett, G., Kushner, J., & Saltonstall, K. (2014). The reproductive biology of *Saccharum spontaneum* L.: Implications for management of this invasive weed in Panama. NeoBiota, 20, 61–79.

Brostoff, W. N. (1989). *Avrainvillea amadelpha* (Codiales, Chlorophyta) from Oahu, Hawaii. Pacific Science, 43, 166–169.

Brown, A. H. D., & Young, A. G. (2000). Genetic diversity in tetraploid populations of the endangered daisy *Rutidosis leptorrhynchoides* and implications for its conservation. Heredity, 85, 122–129.

Charlesworth, D., & Charlesworth, B. (1987). Inbreeding depression and its evolutionary consequences. Annual Review of Ecology, Evolution, and Systematics, 18, 237–268.

Charlesworth, D., & Wright, S. I. (2001). Breeding systems and genome evolution. Current Opinion in Genetics & Development, 11, 685–690.

Clarke, E. (2012). Plant individuality: A solution to the demographer’s dilemma. Biology & Philosophy, 27, 321–361.

Clifton, K. E. (1997). Mass spawning by green algae on coral reefs. Science, 275, 1116–1118.

Clifton, K. E. (2013). The ecological significance of sexual reproduction by tropical green algae. Smithsonian Contributions to the Marine Sciences, 39, 219–228.

Clifton, K. E., & Clifton, L. M. (1999). The phenology of sexual reproduction by green algae (Bryopsidales) on caribbean coral reefs. Journal of Phycology, 35, 24–34.

Comai, L. (2005). The advantages and disadvantages of being polyploid. Nature Reviews Genetics, 6, 836–846.

Combosch, D. J., & Vollmer, S. V. (2011). Population genetics of an ecosystem-defining reef coral *Pocillopora damicornis* in the tropical eastern Pacific. PLOS ONE, 6, e21200.

Conklin, E. J., & Smith, J. E. (2005). Abundance and spread of the invasive red algae, *Kappaphycus* spp., in Kane’ohe Bay, Hawai’i and an experimental assessment of management options. Biological Invasions, 7, 1029–1039.

Cox, T. E., Spalding, H. L., & Foster, M. S. (2017). Spatial and temporal variation of diverse inter-tidal algal assemblages in Southwest O‘ahu. Marine Ecology, 38, e12429.

de Meeûs, T., Prugnolle, F., & Agnew, P. (2007). Asexual reproduction: Genetics and evolutionary aspects. Cellular and Molecular Life Sciences, 64, 1355–1372.

Deveney, M., Rowling, K., Wiltshire, K., Fernandes, M., Collings, G., Tanner, J., & Manning, C. (2008). Caulerpa taxifolia *(M. Vahl) C. Agardh: Environmental risk assessment* (307; South Australian Research and Development Institute Research Report Series, pp. 1–38). South Australian Research and Development Institute.

Doležel, J., Binarová, P., & Lucretti, S. (1989). Analysis of nuclear DNA content in plant cells by flow cytometry. Biologia Plantarum, 31, 113–120.

Doležel, J., Greilhuber, J., & Suda, J. (2007). Estimation of nuclear DNA content in plants using flow cytometry. Nature Protocols, 2, 2233–2244.

Dorken, M. E., & Eckert, C. G. (2001). Severely reduced sexual reproduction in northern populations of a clonal plant, *Decodon verticillatus* (Lythraceae). Journal of Ecology, 89, 339–350.

Doty, M. S. (1961). *Acanthophora*, a possible invader of the marine flora of Hawai’i. Pacific Science, 15, 547–555.

Dueñas, M.-A., Hemming, D. J., Roberts, A., & Diaz-Soltero, H. (2021). The threat of invasive species to IUCN-listed critically endangered species: A systematic review. Global Ecology and Conservation, 26, e01476.

Eckert, C. G., Kalisz, S., Geber, M. A., Sargent, R., Elle, E., Cheptou, P.-O., Goodwillie, C., Johnston, M. O., Kelly, J. K., Moeller, D. A., Porcher, E., Ree, R. H., Vallejo-Marín, M., & Winn, A. A. (2010). Plant mating systems in a changing world. Trends in Ecology & Evolution, 25, 35–43.

Ellstrand, N. C., & Elam, D. R. (1993). Population genetic consequences of small population size: Implications for plant conservation. Annual Review of Ecology, Evolution, and Systematics, 24, 217–242.

Engelen, A. H., Serebryakova, A., Ang, P., Britton-Simmons, K., Mineur, F., Pedersen, M. F., Arenas, F., Fernández, C., Steen, H., Svenson, R., Pavia, H., Toth, G., Viard, F., & Santos, R. (2015). Circumglobal invasion by the brown seaweed *Sargassum muticum*. Oceanography and Marine Biology, 53, 81–126.

Foster, A. D., Spalding, H. L., Cox, T. E., La Valle, F. F., & Philippoff, J. (2019). The invasive green alga *Avrainvillea* sp. Transforms native epifauna and algal communities on a tropical hard substrate reef. Phycological Research, 67, 164–169.

Gill, D. E., Chao Lin, L., Perkins, S. L., & Wolf, J. B. (1995). Genetic mosaicism in plants and clonal animals. Annual Review of Ecology and Systematics, 26, 423–444.

González, A. V., & Santelices, B. (2008). Coalescence and chimerism in *Codium* (Chlorophyta) from central Chile. Phycologia, 47, 468–476.

Govindaraju, D. R. (1988). Relationship between dispersal ability and levels of gene flow in plants. Oikos, 52, 31–35.

Haag, C. R., & Ebert, D. (2004). A new hypothesis to explain geographic parthenogenesis. Annales Zoologici Fennici, 41, 539–544.

Hamrick, J. L., & Godt, M. J. W. (1996). Effects of life history traits on genetic diversity in plant species. Philosophical Transactions of the Royal Society of London. Series B: Biological Sciences, 351, 1291–1298.

Hardy, O. J. (2016). Population genetics of autopolyploids under a mixed mating model and the estimation of selfing rate. Molecular Ecology Resources, 16, 103–117.

Harper, J. L. (1980). Plant demography and ecological theory. Oikos, 35, 244.

Harrison, P. J., Waters, R. E., & Taylor, E. R. J. (1980). A broad spectrum artificial seawater medium for coastal and open ocean phytoplankton. Journal of Phycology, 16, 28–35.

Hay, M. E., Dufy, J. E., Paul, V. J., Renaud, P. E., & Fenical, W. (1990). Specialist herbivores reduce their susceptibility to predation by feeding on the chemically defended seaweed *Avrainvillea longicaulis*. Limnology and Oceanography, 35, 1734–1743.

Hillis-Colinvaux, L. (1984). Systematics of the Siphonales. In D. E. G. Irvine & D. M. John (Eds.), Systematics of the Green Algae (pp. 271–296). Academic Press, Inc.

Kapraun, D. F. (1994). Cytophotometric estimation of nuclear DNA contents in thirteen species of the Caulerpales (Chlorophyta). Cryptogamic Botany, 4, 410–418.

Kapraun, D. F. (2005). Nuclear DNA content estimates in multicellular green, red and brown algae: Phylogenetic considerations. Annals of Botany, 95, 7–44.

Kapraun, D. F., & Nguyen, M. N. (1994). Karyology, nuclear DNA quantification and nucleus-cytoplasmic domain variations in some multinucleate green algae (Siphonocladales, Chlorophyta). Phycologia, 33, 42–52.

Kapraun, D. F., & Shipley, M. J. (1990). Karyology and nuclear DNA quantification in *Bryopsis* (Chlorophyta) from North Carolina, USA. Phycologia, 29, 443–453.

Kittinger, J. N., Bambico, T. M., Minton, D., Miller, A., Mejia, M., Kalei, N., Wong, B., & Glazier, E. W. (2016). Restoring ecosystems, restoring community: Socioeconomic and cultural dimensions of a community-based coral reef restoration project. Regional Environmental Change, 16, 301–313.

Krueger-Hadfield, S. A. (2011). Structure des populations chez l’algue rouge haploid-diploïde Chondrus crispus: Système de reproduction, différeciation génétique et épidémiologie. PhD Dissertation. Université de Pierre et Marie Curie - Sorbonne Université and Pontificia Universidad Católica de Chile.

Krueger-Hadfield, S. A. (2020). What’s ploidy got to do with it? Understanding the evolutionary ecology of macroalgal invasions necessitates incorporating life cycle complexity. Evolutionary Applications, 13, 486–499.

Krueger-Hadfield, S. A. (2024) Let’s talk about sex: why reproductive systems matter for understanding algae. Journal of Phycology, doi: 10.1111/jpy.13462.

Krueger-Hadfield, S. A., Guillemin, M.-L., Destombe, C., Valero, M., & Stoeckel, S. (2021). Exploring the genetic consequences of clonality in haplodiplontic taxa. Journal of Heredity, 112, 92–107.

Krueger-Hadfield, S. A., Kollars, N. M., Byers, J. E., Greig, T. W., Hammann, M., Murray, D. C., Murren, C. J., Strand, A. E., Terada, R., Weinberger, F., & Sotka, E. E. (2016). Invasion of novel habitats uncouples haplo-diplontic life cycles. Molecular Ecology, 25, 3801–3816.

Krueger-Hadfield, S. A., Kollars, N. M., Strand, A. E., Byers, J. E., Shainker, S. J., Terada, R., Greig, T. W., Hammann, M., Murray, D. C., Weinberger, F., & Sotka, E. E. (2017). Genetic identification of source and likely vector of a widespread marine invader. Ecology and Evolution, 7, 4432–4447.

Krueger-Hadfield, S. A., Oetterer, A. P., Lees, L. E., Hoffman, J. M., Sotka, E. E., & Murren, C. J. (2023). Phenology and thallus size in a non-native population of *Gracilaria vermiculophylla*. Journal of Phycology, 59, 926–938.

Krueger-Hadfield, S. A., Roze, D., Mauger, S., & Valero, M. (2013). Intergametophytic selfing and microgeographic genetic structure shape populations of the intertidal red seaweed *Chondrus crispus*. Molecular Ecology, 22, 3242–3260.

Lagourgue, L., Rousseau, F., Zubia, M., & Payri, C. E. (2023). Diversity of the genus *Avrainvillea* (Dichotomosiphonaceae, Chlorophyta): New insights and eight new species. European Journal of Phycology, 58, 399–426.

Lane, J. E., Forrest, M. N. K., & Willis, C. K. R. (2011). Anthropogenic influences on natural animal mating systems. Animal Behaviour, 81, 909–917.

Langston, R. C., & Spalding, H. L. (2017). A survey of fishes associated with Hawaiian deep-water *Halimeda kanaloana* (Bryopsidales: Halimedaceae) and *Avrainvillea* sp. (Bryopsidales: Udoteaceae) meadows. PeerJ, 5, e3307.

Littler, D. S., & Littler, M. M. (1992). Systematics of *Avrainvillea* (Bryopsidales, Chlorophyta) in the tropical western Atlantic. Phycologia, 31, 375–418.

Littler, M. M., Littler, D. S., & Brooks, B. L. (2004). Extraordinary mound-building forms of *Avrainvillea* (Bryopsidales, Chlorophyta): Their experimental taxonomy, comparative functional morphology, and ecological strategies. Atoll Research Bulletin, 515, 1–26.

Littler, M. M., Littler, D. S., & Brooks, B. L. (2005). Extraordinary mound building *Avrainvillea* (Chlorophyta): The largest tropical marine plants. Coral Reefs, 24, 555.

Longenecker, K., Bolick, H., & Kawamoto, R. (2011). *Macrofaunal invertebrate communities on Hawaii’s shallow coral-reef flats: Changes associated with the removal of an invasive alien alga* (Bishop Museum Technical Report 54; Hawaii Biological Survey, pp. 1–51). Bishop Museum.

Luttikhuizen, P. C., Stift, M., Kuperus, P., & Van Tienderen, P. H. (2007). Genetic diversity in diploid vs. Tetraploid *Rorippa amphibia* (Brassicaceae). Molecular Ecology, 16, 3544– 3553.

Lynch, M. (1984). Destabilizing hybridization, general-purpose genotypes and geographic parthenogenesis. The Quarterly Review of Biology, 59, 257–290.

Magalhães, W. F., & Bailey-Brock, J. H. (2014). Polychaete assemblages associated with the invasive green alga *Avrainvillea amadelpha* and surrounding bare sediment patches in Hawaii. Memoirs of Museum Victoria, 71, 161–168.

Marshall, D. R., & Weir, B. (1979). Maintenance of genetic variation in apomictic plant populations. I. Single locus models. Heredity, 42, 159–172.

Meirmans, P. G. (2020). GENODIVE version 3.0: Easy-to-use software for the analysis of genetic data of diploids and polyploids. Molecular Ecology Resources, 20, 1126–1131.

Meirmans, P. G., & Liu, S. (2018). Analysis of molecular variance (AMOVA) for autopolyploids. Frontiers in Ecology and Evolution, 6, 66.

Meirmans, P. G., & Van Tienderen, P. H. (2013). The effects of inheritance in tetraploids on genetic diversity and population divergence. Heredity, 110, 131–137.

Melnyk, C. W., & Meyerowitz, E. M. (2015). Plant grafting. Current Biology, 25, R183–R188.

Moody, M. E., Mueller, L. D., & Soltis, D. E. (1993). Genetic variation and random drift in autotetraploid populations. Genetics, 134, 649–657.

Mudge, K., Janick, J., Scofield, S., & Goldschmidt, E. E. (2009). A history of grafting. Horticultural Reviews, 35, 437–493.

Nei, M. (1978). Estimation of average heterozygosity and genetic distance from a small number of individuals. Genetics, 89, 583–590.

Olsen, K. C., Ryan, W. H., Winn, A. A., Kosman, E. T., Moscoso, J. A., Krueger-Hadfield, S. A., Burgess, S. C., Carlon, D. B., Grosberg, R. K., Kalisz, S., & Levitan, D. R. (2020). Inbreeding shapes the evolution of marine invertebrates. Evolution, 74, 871–882.

Olsen-Stojkovich, J. (1979). Revision of the pantropical algal genus Avrainvillea Decaisne (Codiales, Codiaceae) M.S. Thesis. University of Guam.

Olsen-Stojkovich, J. (1985). A systematic study of the genus *Avrainvillea* Decaisne (Chlorophyta, Udoteaceae). Nova Hedwigia, 41, 1–68.

Orive, M. E., & Krueger-Hadfield, S. A. (2021). Sex and asex: A clonal lexicon. Journal of Heredity, 112, 1–8.

Otto, S. P. (2007). The evolutionary consequences of polyploidy. Cell, 131, 452–462.

Otto, S. P., & Marks, J. C. (1996). Mating systems and the evolutionary transition between haploidy and diploidy. Biological Journal of the Linnean Society, 57, 197–218.

Pannell, J. R. (2015). Evolution of the mating system in colonizing plants. Molecular Ecology, 24, 2018–2037.

Pannell, J. R., Auld, J. R., Brandvain, Y., Burd, M., Busch, J. W., Cheptou, P.-O., Conner, J. K., Goldberg, E. E., Grant, A.-G., Grossenbacher, D. L., Hovick, S. M., Igic, B., Kalisz, S., Petanidou, T., Randle, A. M., de Casas, R. R., Pauw, A., Vamosi, J. C., & Winn, A. A. (2015). The scope of Baker’s law. New Phytologist, 208, 656–667.

Parisod, C., Holderegger, R., & Brochmann, C. (2010). Evolutionary consequences of autopolyploidy. New Phytologist, 186, 5–17.

Peck, J. R., Yearsley, J. M., & Waxman, D. (1998). Explaining the geographic distributions of sexual and asexual populations. Nature, 391, 889–892.

Petitpierre, B., Kueffer, C., Broennimann, O., Randin, C., Daehler, C., & Guisan, A. (2012). Climatic niche shifts are rare among terrestrial plant invaders. Science, 335, 1344–1348.

Peyton, K. A. (2009). Aquatic invasive species impacts in Hawaiian soft sediment habitats [Ph.D.]. The University of Hawaiʻi at Mānoa.

Preston, R., Blomster, J., Schagerström, E., & Seppä, P. (2022). Clonality, polyploidy and spatial population structure in Baltic Sea *Fucus vesiculosus*. Ecology and Evolution, 12, e9336.

Quarin, C. L., Espinoza, F., Martinez, E. J., Pessino, S. C., & Bovo, O. A. (2001). A rise of ploidy level induces the expression of apomixis in *Paspalum notatum*. Sexual Plant Reproduction, 13, 243–249.

R Core Team. (2018). R: A Language and Environment for Statistical Computing (3.5.2). R Foundation for Statistical Computing. https://www.R-project.org/

Reichel, K., Masson, J.-P., Malrieu, F., Arnaud-Haond, S., & Stoeckel, S. (2016). Rare sex or out of reach equilibrium? The dynamics of *F_IS_* in partially clonal organisms. BMC Genetics, 17, 76.

Ronfort, J., Jenczewski, E., Bataillon, T., & Rousset, F. (1998). Analysis of population structure in autotetraploid species. Genetics, 150, 921–930.

Rushworth, C. A., Wagner, M. R., Mitchell-Olds, T., & Anderson, J. T. (2022). The Boechera model system for evolutionary ecology. American Journal of Botany, 109, 1939–1961.

Santelices, B., Aedo, D., Hormazábal, M., & Flores, V. (2003). Field testing of inter-and intraspecific coalescence among mid-intertidal red algae. Marine Ecology Progress Series, 250, 91–103.

Santelices, B., Correa, J. A., Aedo, D., Flores, V., Hormazábal, M., & Sánchez, P. (1999). Convergent biological processes in coalescing Rhodophyta. Journal of Phycology, 35, 1127–1149.

Schaffelke, B., Smith, J. E., & Hewitt, C. L. (2006). Introduced macroalgae – a growing concern. Journal of Applied Phycology, 18, 529–541.

Scrosati, R. (2002). An updated definition of genet applicable to clonal seaweeds, bryophytes, and vascular plants. Basic and Applied Ecology, 3, 97–99.

Smith, J. E., Hunter, C. L., Conklin, E. J., Most, R., Sauvage, T., Squair, C., & Smith, C. M. (2004). Ecology of the invasive red alga *Gracilaria salicornia* (Rhodophyta) on O’ahu, Hawai’i. Pacific Science, 58, 325–343.

Smith, J. E., Hunter, C. L., & Smith, C. M. (2002). Distribution and reproductive characteristics of nonindigenous and invasive marine algae in the Hawaiian Islands. Pacific Science, 56, 299–315.

Soltis, D. E., & Soltis, P. S. (1999). Polyploidy: Recurrent formation and genome evolution. Trends in Ecology & Evolution, 14, 348–352.

Sotka, E. E., Baumgardner, A. W., Bippus, P. M., Destombe, C., Duermit, E. A., Endo, H., Flanagan, B. A., Kamiya, M., Lees, L. E., Murren, C. J., Nakaoka, M., Shainker, S. J., Strand, A. E., Terada, R., Valero, M., Weinberger, F., & Krueger-Hadfield, S. A. (2018). Combining niche shift and population genetic analyses predicts rapid phenotypic evolution during invasion. Evolutionary Applications, 11, 781–793.

Sousa, F., Neiva, J., Martins, N., Jacinto, R., Anderson, L., Raimondi, P. T., Serrão, E. A., & Pearson, G. A. (2019). Increased evolutionary rates and conserved transcriptional response following allopolyploidization in brown algae. Evolution, 73, 59–72.

Spalding, H. L. (2012). Ecology of mesophotic macroalgae and Halimeda kanaloana meadows in the main Hawaiian islands. Ph.D. Dissertation. The University of Hawaiʻi at Mānoa.

Spalding, H. L., Copus, J. M., Bowen, B. W., Kosaki, R. K., Longenecker, K., Montgomery, A. D., Padilla-Gamiño, J. L., Parrish, F. A., Roth, M. S., Rowley, S. J., Toonen, R. J., & Pyle, R. L. (2019). The Hawaiian Archipelago. In Y. Loya, K. A. Puglise, & T. C. L. Bridge (Eds.), Mesophotic Coral Ecosystems (pp. 445–464). Springer International Publishing.

Stoeckel, S., Arnaud-Haond, S., & Krueger-Hadfield, S. A. (2021). The combined effect of haplodiplonty and partial clonality on genotypic and genetic diversity in a finite mutating population. Journal of Heredity, 112, 78–91.

Stoeckel, S., Becheler, R., Bocharova, E., & Barloy, D. (2024). GENAPOPOP 1.0: A user-friendly software to analyse genetic diversity and structure from partially clonal and selfed autopolyploid organisms. Molecular Ecology Resources, 00, 1–11.

Stoeckel, S., & Masson, J.-P. (2014). The exact distributions of *F_IS_* under partial asexuality in small finite populations with mutation. PLOS ONE, 9, e85228.

te Beest, M., Le Roux, J. J., Richardson, D. M., Brysting, A. K., Suda, J., Kubesova, M., & Pysek, P. (2012). The more the better? The role of polyploidy in facilitating plant invasions. Annals of Botany, 109, 19–45.

Tuomi, J., & Vuorisalo, T. (1989). What are the units of selection in modular organisms? Oikos, 54, 227.

Van De Verg, S. E., & Smith, C. M. (2022). Protocol to control the invasive alga *Avrainvillea lacerata* in a shallow Hawaiian reef flat. Applications in Plant Sciences, 10, e11490.

Vandel, A. (1928). La parthénogénèse geographique. Contribution à l’étude biologique et cytologique de la parthénogénèse naturelle. Bulletin Biologique de La France et de La Belgique, 62, 164–281.

Varela-Álvarez, E., Gómez Garreta, A., Rull Lluch, J., Salvador Soler, N., Serrao, E. A., & Siguán, M. A. R. (2012). Mediterranean species of *Caulerpa* are polyploid with smaller genomes in the invasive ones. PLoS ONE, 7, e47728.

Varela-Álvarez, E., Loureiro, J., Paulino, C., & Serrão, E. A. (2018). Polyploid lineages in the genus *Porphyra*. Scientific Reports, 8, 8696.

Veazey, L., Williams, O., Wade, R., Toonen, R., & Spalding, H. L. (2019). Present-day distribution and potential spread of the invasive green alga *Avrainvillea amadelpha* around the main Hawaiian Islands. Frontiers in Marine Science, 6, 402.

Verbruggen, H., Ashworth, M., LoDuca, S. T., Vlaeminck, C., Cocquyt, E., Sauvage, T., Zechman, F. W., Littler, D. S., Littler, M. M., Leliaert, F., & De Clerck, O. (2009). A multi-locus time-calibrated phylogeny of the siphonous green algae. Molecular Phylogenetics and Evolution, 50, 642–653.

Voisin, M., Engel, C. R., & Viard, F. (2005). Differential shuffling of native genetic diversity across introduced regions in a brown alga: Aquaculture vs. maritime traffic effects. Proceedings of the National Academy of Sciences, 102, 5432–5437.

Wade, R. M. (2019). An alvigorous sea slug as a novel sampling tool and its implications for algal diversity, herbivore ecology, and invasive species tracking. Ph.D. Dissertation. The University of Hawaiʻi at Mānoa.

Wade, R. M., Spalding, H. L., Peyton, K. A., Foster, K., Sauvage, T., Ross, M., & Sherwood, A. R. (2018). A new record of *Avrainvillea* cf. *Erecta* (Berkeley) A. Gepp & E. S. Gepp (Bryopsidales, Chlorophyta) from urbanized estuaries in the Hawaiian Islands. Biodiversity Data Journal, 6, e21617.

Wai, M. K., Nyunt, T., Kyaw, S. P. P., & Soe-Htun, U. (2009). The morphology and distribution of the genus *Avrainvillea* (Bryopsidales, Chlorophyta) from Myanmar. Journal of the Myanmar Academy of Arts and Science, 7, 199–211.

Weir, B. (1996). Genetic data analysis II: Methods for discrete population genetic data. Sinauer Associates Inc.

Weir, B. S., & Cockerham, C. C. (1984). Estimating F-statistics for the analysis of population structure. Evolution, 38, 1358–1370.

Whitehead, M. R., Lanfear, R., Mitchell, R. J., & Karron, J. D. (2018). Plant mating systems often vary widely among populations. Frontiers in Ecology and Evolution, 6, 38.

Williams, S. L., & Schroeder, S. L. (2004). Eradication of the invasive seaweed *Caulerpa taxifolia* by chlorine bleach. Marine Ecology Progress Series, 272, 69–76.

Williams, S. L., & Smith, J. E. (2007). A global review of the distribution, taxonomy, and impacts of introduced seaweeds. Annual Review of Ecology, Evolution, and Systematics, 38, 327–359.

Williams, T. M., Krueger-Hadfield, S. A., Hill-Spanik, K. M., Kosaki, R. K., Stoeckel, S., & Spalding, H. L. (2024). The reproductive system of the cryptogenic alga *Chondria tumulosa* (Florideophyceae) at Manawai, Papahānaumokuākea Marine National Monument. Phycologia, 63, 36–44.

Wilson, J. R. U., Dormontt, E. E., Prentis, P. J., Lowe, A. J., & Richardson, D. M. (2009). Something in the way you move: Dispersal pathways affect invasion success. Trends in Ecology & Evolution, 24, 136–144.

Zamora, D. L., Thill, D. C., & Eplee, R. E. (1989). An Eradication Plan for Plant Invasions. Weed Technology, 3, 2–12.

